# QTL for Heat-Induced Stomatal Anatomy Underpin Gas Exchange Variation in Field-Grown Wheat

**DOI:** 10.64898/2025.12.16.694723

**Authors:** Edward Chaplin, Emi Tanaka, Andrew Merchant, Beata Sznajder, Richard Trethowan, William Salter

## Abstract

Stomata are central to leaf gas exchange, governing carbon uptake, water loss, and ultimately, crop performance. However, the contribution of integrated stomatal anatomy and physiology to wheat heat tolerance remains poorly understood, particularly under realistic field conditions and across divers germplasm. This study explored the role of stomatal anatomical and physiological traits in shaping wheat responses to heat stress.

Across two years of multi-environment field trials encompassing 200 genotypes in season 1 and 50 genotypes in season 2, we examined stomatal conductance (*g*ₛ), anatomical traits including stomatal size and density, and the stomatal conductance operating efficiency (*g_se_*) across leaf surfaces, along with grain yield. Timely and delayed sowing treatments were used to expose key developmental stages (anthesis) to contrasting temperature regimes.

Early sowing supported higher *g_s_* and *g_se_*, while delayed sowing impaired stomatal function despite similar theoretical anatomical capacity (*g_smax_*), revealing a decoupling of structural potential and physiological performance under stress. The adaxial surface consistently exhibited higher *g_s_*, stomatal density, and *g_smax_* than the abaxial surface, highlighting its dominant role in leaf gas exchange. Later sowing induced plastic shifts in anatomy, including smaller, denser stomata, particularly on the adaxial surface, suggesting an adaptive response to thermal stress.

Significant genotypic variation was observed for *g_s_, g_se_, g_smax_*, and stomatal anatomical traits, with moderate heritability indicating genetic control. 125 putative QTL were identified for multiple stomatal traits across environments, including several stable loci on chromosomes 2B and 5B, and numerous closely clustered QTL for anatomical traits on chromosome 7B, highlighting key genomic regions underlying stomatal anatomy. In contrast, QTL for *g_s_* and *g_se_* were fewer and season-specific, highlighting the environmentally sensitive nature of physiological stomatal regulation. 42 of the QTL identified were consistent with previously reported QTL for stomatal traits in wheat.

Together, these findings elevate the role of stomatal traits, supporting an integrated breeding strategy that combines selection for favourable stomatal anatomy with efficiency physiological regulation. Incorporating traits like *g_se_* into selection frameworks may enhance yield stability and resilience in heat-prone environments, advancing the development of climate-resilient wheat ideotypes.

**Scope:** This manuscript examines how stomatal anatomical and physiological traits integrate to shape wheat responses to heat stress under field conditions, addressing a central challenge in understanding the roles of stomata in a warming climate. Using multi-environment field trials across two growing seasons and encompassing 200 wheat genotypes, we quantify stomatal conductance (*g_s_*), anatomical traits, and stomatal operating efficiency (*g_se_*), across adaxial and abaxial leaf surfaces, and investigate whether these are under genetic control.

The study provides mechanistic insight into the dynamic regulation of stomatal function by demonstrating a decoupling between anatomical capacity and physiological performance under heat stress, alongside plastic shifts in stomatal traits across sowing times. The identification of 125 candidate QTLs, including numerous stable and co-localised QTL across seasons, supports their incorporation into breeding programs aimed at enhancing resilience and yield stability.

This work aligns with the Research Topic by elucidating mechanistic underpinnings of stomatal conductance regulation, bridging stomatal biology with applied crop improvement strategies. This work is critical to improving understanding of plant water relations and carbon uptake under future climate scenarios to ensure food security in a changing climate.

## Background

Heat stress is a major constraint to global crop productivity, particularly in wheat (*Triticum aestivum* L.), the world’s most widely cultivated cereal (Erenstein et al., 2022). As a staple food for over one-third of the global population (IRDC, 2010) and a cornerstone of Australian agriculture (FAOSTAT, 2024), wheat is especially vulnerable to rising temperatures. Projected global temperature increases of 1.8–5.7°C by 2100 (OECD, 2012) will intensify the frequency, severity, and duration of heatwaves, particularly in major wheat-growing regions such as the Australian Grain Belt (Anwar et al., 2013; Collins et al., 2024; Perkins-Kirkpatrick and Lewis, 2020). Extreme heat events during critical developmental stages, such as anthesis, cause substantial yield losses and threaten global food security (Pequeno et al., 2021; Ullah et al., 2022; Yashavanthakumar et al., 2021).

Stomata play a central role in balancing photosynthetic carbon assimilation and evaporative cooling (Lawson and Blatt, 2014; Xie et al., 2022). Under heat stress, this balance is disrupted by both stomatal and non-stomatal limitations including damage to photosystem II (PSII), reduced Rubisco carboxylation, impaired regeneration of ribulose-1,5-bisphosphate (RuBP) (Chaves and Oliveira, 2004; Chaves et al., 2009), and restricted CO₂ uptake (Eisenhut et al., 2019; Hodges et al., 2016). The consequences for wheat yield are substantial, with every 1°C rise in temperature reducing wheat yields by 6-10% (Asseng et al., 2015; Helman and Bonfil, 2022; Liu et al., 2016; Shew et al., 2020) and Australian yields projected to fall by 10-30% by 2050 (Liu et al., 2016; Potgieter et al., 2013; Zeleke, 2021). Nevertheless, opportunities exist to enhance yield resilience by targeting physiological traits and processes that sustain photosynthetic efficiency under heat stress.

Stomatal conductance (*g*ₛ), the rate at which CO₂ and water vapour diffuse through the stomata (Wall et al., 2022), directly influences photosynthesis and canopy cooling, but also governs water loss (Roche, 2015; Wong et al., 1979). While *g*ₛ responds almost instantaneously to environmental cues through physiological adjustments, its upper and lower bounds are constrained by longer-term anatomical traits, such as stomatal size and density (SD; number of stomata per unit leaf area) (Lawson and Leakey, 2024). Wheat shows wide genotypic variation in these traits (Faralli et al., 2024; Lawson and Matthews, 2020; Stevens et al., 2021). As an amphistomatous species, wheat typically exhibits higher adaxial SD, and in turn is the key driver of total leaf *g*ₛ (Hõrak, 2025; McAusland et al., 2021; Muir, 2015; Wall et al., 2022). SD is the primary determinant of the maximum potential anatomical stomatal conductance (*g_smax_*), setting the upper physiological limit for gas exchange (Franks et al., 2009; Franks and Beerling, 2009). Integrating understanding of these anatomical constraints with operating stomatal conductance (*g_sop_*), provides critical insights into gas exchange efficiency (*g_se_*; *g_se_ = g_sop_/g_smax_*) and heat tolerance mechanisms, allowing identification of optimal stomatal traits conferring enhanced heat tolerance (Busch et al., 2024; Haworth et al., 2021). However, adaxial stomata often realise less of their anatomical *g_smax_* and show greater sensitivity to closure than abaxial stomata, highlighting surface-specific asymmetry in stomatal behaviour (Driesen et al., 2023; Fanourakis et al., 2015; Tulva et al., 2025).

Although responses vary, most studies report reduced *g_s_* under heat stress as a water-conserving response that limits transpiration (Balla et al., 2019; Pooja and Munjal, 2019; Redhu et al., 2025) but also restricts CO_2_ uptake (Monneveux et al., 2003). Conversely, several studies show increased *g_s_* under heat when water is not limiting, suggesting a strategy to enhance evaporative cooling and maintain photosynthetic function (Abdelhakim et al., 2021; Djanaguiraman et al., 2020; Mirosavljević et al., 2021), or reflecting the inherent heat-tolerance of certain germplasm (Abdelhakim et al., 2021; Pinto et al., 2025; Ramya et al., 2016; Sharma et al., 2014, 2014). These divergent patterns highlight the context-dependency of stomatal behaviour, shaped by interactions between water availability, evaporative demand and genotypic capacity for thermal regulation. Large genotypic variation in *g*_s_ (Mahdavi et al., 2021; Pooja and Munjal, 2019), heritability estimates of up to 73%, and strong associations between stomatal traits, photosynthetic assimilation, and yield (Li et al., 2021; McAusland et al., 2021; Pinto et al., 2025) highlight significant potential for genetic improvement (Rebetzke et al., 2003, 2001; Romena and Farshadfar, 2019).

Few studies have investigated stomatal anatomical responses to heat stress at breeding-relevant scales, although reduced stomatal pore size and increased SD have been reported in studies with fewer genotypes (Fan et al., 2022; Kapadiya et al., 2017; Li et al., 2023). Smaller, denser stomata can open and close more rapidly, facilitating tighter control over gas exchange and evaporative cooling (Drake et al., 2013; McAusland et al., 2016). As with *g_s_*, anatomical responses to heat stress are often stronger on the adaxial surface (Bramley et al., 2022; Pinto et al., 2025). Substantial genotypic variation in anatomical traits indicate potential for selection (Condon et al., 2007; McAusland et al., 2021; Pinto et al., 2025).

Additionally, high heritability and QTLs have been reported for stomatal anatomical traits, some with pleiotropic effects on yield (Liu et al., 2025; Romena and Farshadfar, 2019; Shahinnia et al., 2016; Wang et al., 2016, 2015). Yet, despite the role of stomatal anatomical traits in shaping kinetic responses and heat tolerance, they remain underexplored in breeding contexts.

Going forwards, a more integrated approach is needed to link anatomical potential with in season physiological performance, considering their interplay alongside environmental conditions and agronomic outcomes like yield (Yan, 2021). Incorporating coupled measurements of *g_s_* and anatomy provides deeper insights into gas exchange efficiency, uncovering scenarios where there is disconnect between anatomical capacity and realised function, for example, where *g_smax_* is maintained or increased but *g_s_* is reduced (Dow et al., 2014). Such integration also supports identification of shared genetic controls across traits and mitigation of trade-offs in water-use-efficiency (Dwivedi et al., 2021).

Despite their importance in carbon assimilation and WUE (Roche, 2015), progress in scaling precise stomatal measurements to breeding contexts has been limited by methodological constraints. Infrared gas analysers (IRGAs) to measure *g_s_* are accurate but slow and labour-intensive (Guizani et al., 2023; Wang et al., 2024), while nail polish imprints to assess stomatal anatomy lack field-suitable scalability (Millstead et al., 2020; Wall et al., 2023, 2022). Recent advances in high-throughput phenotyping using handheld porometers (e.g., LI-COR LI-600) (McAusland et al., 2023; Reynolds et al., 2020) and handheld digital microscopes (Liang et al., 2022; Pathoumthong et al., 2023; Sun et al., 2023, 2021) enable efficient, non-destructive, in-situ measurements of *g_s_* and anatomy, while integration with deep learning has greatly expanded the scalability of stomatal phenotyping (Chaplin et al., 2025a; Gibbs and Burgess, 2024).

This study examines the effects of heat stress on *g_s_*, stomatal anatomy, and key physiological indicators such as *g_se_* and *g_smax_* in a diverse panel of 200 wheat genotypes over two seasons, as well as assessing grain yield. We conducted two independent field trials over consecutive seasons: (1) a 2023 trial comparing 200 genotypes and (2) a 2024 trial assessing 50 selected genotypes, both with two time of sowing (TOS) treatments. We hypothesised delayed sowing would reduce *g_sop_*, accompanied by an increase in SD and a reduction in stomatal size. Additionally, we expected significant genotypic variation in stomatal traits which would support the identification of markers and QTLs associated with these traits for future crop improvement efforts. This study fills key gaps by linking stomatal physiology and anatomy – traits often examined in isolation – to provide a holistic view of heat stress responses.

Adopting novel high-throughput phenotyping tools to screen 200 diverse wheat genotypes across two seasons in large, replicated field trials, it offers an unprecedented opportunity to evaluate how stomatal traits interact under heat stress. By combining extensive physiological and anatomical data, our findings aim to inform breeding strategies for climate-resilient wheat.

## Methods

### Plant Material, Germplasm G Experimental Conditions

Field plots were established at the University of Sydney I.A. Watson Grains Research Centre in Narrabri, NSW Australia (30.2743°S, 149.8093°E). Two field trial experiments on wheat (*Triticum aestivum* L.) were carried out over two years (2023 & 2024) to investigate the effect of heat stress, simulated through timely and delayed sowing treatments. In 2023 (Season 1; S1), 200 genotypes (Table 1) were established with two planting dates; ‘TOS 1’ and ‘TOS 2’. A randomised complete block design with two field replicates nested within a larger training population was used. In 2024 (Season 2; S2), 50 genotypes (Table 2), all common to those grown in S1 selected based upon diversity in heat tolerance of physiological traits, were established with two planting dates; TOS 1 and TOS 2. In both years, two field replicate plots arranged in randomised blocks were sown for each TOS.

**Table 1:**
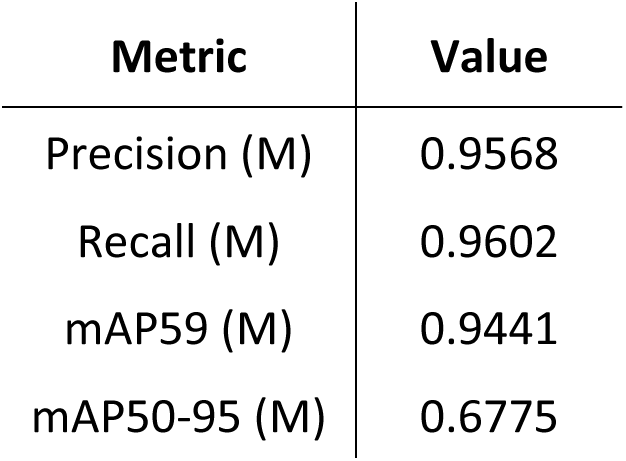
Performance metrics for YOLOv8-M model. Metrics used were: Precision, Recall, mAP50 (mean average precision at Intersection over Union = 0.5) and mAP50-95 (mean average precision at Intersection over Union from 0.5 to 0.95).

| Metric | Value |
| --- | --- |
| Precision (M) | 0.9568 |
| Recall (M) | 0.9602 |
| mAP59 (M) | 0.9441 |
| mAP50-95 (M) | 0.6775 |

**Table 2:** Linear mixed model ANOVA p values for TOS, Variety and Surface on traits of wheat from 2023 field data.

| Trait | *p*<br>values |  |  |  |  |  |  |
| --- | --- | --- | --- | --- | --- | --- | --- |
|  | TOS | Variety | Surface | TOS x<br>Variety | TOS x<br>Surface | Variety x<br>Surface | TOS x Variety x<br>Surface |
| **Stomatal Conductance** | <0.001 | 0.505 | <0.001 | 0.897 | <0.001 | 0.854 | 0.579 |
| **Guard Cell Width** | <0.001 | <0.001 | <0.001 | 0.545 | 0.383 | 0.183 | 0.479 |
| **Guard Cell Length** | <0.001 | <0.001 | 0.322 | 0.248 | 0.002 | 0.193 | 0.470 |
| **Stomatal Area** | <0.001 | <0.001 | 0.211 | 0.260 | 0.002 | 0.201 | 0.505 |
| **Stomatal Density** | <0.001 | <0.001 | <0.001 | 0.717 | 0.174 | 0.010 | 0.491 |
| ***g<sub>smax</sub>*** | 0.490 | 0.0248 | <0.001 | 0.280 | 0.527 | 0.015 | 0.310 |
| ***g<sub>se</sub>*** | <0.001 | 0.288 | <0.001 | 0.989 | <0.001 | 0.571 | 0.650 |
| **Yield** | <0.001 | <0.001 | - | <0.001 | - | - | - |

The diverse germplasm comprised CIMMYT (International Maize and Wheat Improvement Center) and ICARDA (International Centre for Agricultural Research in the Dry Areas) heat tolerant materials imported through the CIMMYT Australia ICARDA Germplasm Evaluation (CAIGE) program. Lines include materials from SATYN (Stress Adapted Trait Yield Nursery), EDPIE (Elite Diversity International Experiment), ESWYT (Elite Selection Wheat Yield Trial), SAWYT (Semi-Arid Wheat Yield Trial), HTWYT (High Temperature Wheat Yield Trials) and the University of Sydney including recombinants from 3 cycles of genomic selection for heat tolerance (including progeny derived from crosses among CIMMYT and ICARDA materials).

Plants were sown in 12 m^2^ plots (2 × 6 m) with five planting rows at a planting density of 100 plants/m^2.^ Plots were subsequently trimmed 8 m^2^ (2 × 4 m) before harvest. The distance between rows in each plot (row spacing) was approximately 23.5 cm and the distance between each plot was 63 cm. The predominant soil type at the field site is a black vertosol cracking clay with high water retention. For both trials, minimum tillage was used to maintain soil integrity and moisture. Field traffic was controlled as much as possible with allocated roads and pathways and GPS guidance on all machinery. Fertiliser application was kept consistent across both treatments and both trials (Urea [46% N] 100 kg/ha and Cotton Sustain [5% N, 10% P, 21% K, 1% Z] 80 kg/ha pre-planting). The experimental sites were fallowed over the summer months and rotated with a legume crop during alternate years to minimise disease outbreak and maintain soil integrity.

In S1, TOS 1 was planted on 30.05.2023 and harvested on 31.10.2023 while TOS 2 was planted on 19.07.2023 and harvested on 13.12.2023. At planting, average soil moisture was 18.8% at 10cm, 19.5% at 30cm, 17.3% at 50cm and 19.9% at 100cm at TOS 1 and 31.6%, 36.8%, 34.1%, and 36.6% at TOS 2 at 10cm, 30cm, 50cm and 100cm, respectively. Irrigation was used to limit the confounding effects of moisture stress, particularly in later date of sowing. TOS 1 plants received 150 mm of rainfall and irrigation while TOS 2 received 227.9 mm of total moisture from rainfall and irrigation. The mean daily minimum and maximum temperature across the growing season were 6.1 °C and 22.8 °C for TOS 1, respectively, and 9.7 °C and 27.2 °C for TOS 2, respectively.

In S2, TOS 1 was planted on 22.05.2024 and harvested on 06.11.2024, and TOS 2 was planted on 23.07.2024 and harvested on 27.11.2024. At planting, average soil moisture was 18.2% at 10cm, 15.6% at 30cm, 14.6% at 50cm and 11.1% at 100cm at TOS 1 and 28.0%, 31.4%, 31.7% and 31.0% at TOS 2 at 10cm, 30cm, 50cm and 100cm, respectively. Irrigation was used to limit the confounding effects of moisture stress, particularly in later date of sowing. The total moisture from rainfall and irrigation for TOS 1 and TOS 2 was 278.4 mm and 160.4 mm, respectively. The mean daily minimum and maximum temperature across the growing season were 6.4 °C and 21.2 °C for TOS 1, respectively, and 8.8 °C and 25.1 °C for TOS 2, respectively.

### Field Measurements

In S1 and S2, measurements of stomatal physiology and anatomy were taken from fully expanded flag leaves at anthesis (Zadok stage 48-68) and chosen using a systematic randomised sampling technique with representative plants selected from the middle three planting rows of each plot, at least 50 cm from the end of each plot. In all seasons, data were collected on consecutive days and all measurements were taken between 09:30 and 14:00 to minimise diurnal effects on stomatal conductance and rates of photosynthesis. In S1, two leaves were sampled from each replicate plot per TOS (n=4 per genotype per TOS). In S2, three leaves were sampled from each replicate plot per TOS (n=6 per genotype per TOS).

### Stomatal Conductance

In S1 and S2, a LI-COR LI-600 porometer/fluorometer (Li-COR Inc., Nebraska, USA) was used to measure *g_s_* of both leaf surfaces (adaxial and abaxial) (Figure S1). The settings were as follows: flow rate: 150 µmol s^-1^, phase length: 300 ms, ramp amount: 25%, leaf absorptance: 0.8, fraction abs PSII: 0.5, actinic modulation rate: 500 Hz, and integrated modulation intensity: 6.67 µmol m^-2^ s^-1^, stability: medium. Measurements were collected from the middle portion of the leaf blade from the flag leaf of the main tiller in full sunlight, as described by Chaplin et al., (2025).

### Stomatal Anatomy Image Capture

In S1 and S2, in-field microscopy imaging of the abaxial and adaxial surfaces of the same leaf was carried out. In S1, a 200x magnification handheld USB microscope was used (Dino-Lite 5MP, AM7515MT2A; Dino-Lite, AnMo Electronics Corporation, Taiwan) and in S2, a 400x magnification model was used (Dino-Lite 5MP, AM7515MT4A; Dino-Lite, AnMo Electronics Corporation, Taiwan). In both seasons, the microscope was configured with LED brightness = 5, LED axial light = 0 and resolution = 2592×1944.

To aid focussing of the microscope in the field, a custom designed 3D printed leaf clip with a smaller aperture than the cap provided with the microscope was used. Black electrical tape was wrapped around the clear plastic part of the microscope shaft to ensure consistent lighting conditions and prevent external light from hitting the leaf surface. The microscope was connected by USB cable to a Windows computer (Dell Latitude 7230 Rugged Extreme Tablet). Each leaf was imaged by clamping the leaf into the leaf clip, on the same portion of leaf measured with the porometer. Fine focusing on the leaf surface was achieved by adjusting pressure on the trigger with the leaf clamped in the leaf clip. In S2, following image capture, leaves were then excised and placed in an envelope for subsequent nutrient component analysis. Images were saved using a custom image capture app, FieldDino, which was built with Python and PyQt5, and using the Dino-Lite SDK. The app and guidance on its installation are available on the Github repository (https://github.com/williamtsalter/FieldDinoMicroscopy). As described by Chaplin et al., (2025), it incorporates key elements of field data collection:

1. Pre-filled image names based on an existing spreadsheet to ensure each image matches the plot in which it is collected
2. Clear video feed with recordable and adjustable microscope parameters
3. Easy to use buttons and image deletion

### Yield and Yield Components

At maturity, plots were machine-harvested and grain yield was recorded and expressed on a per-hectare basis. In season 2, subsamples of cleaned grain were analysed for quality traits. Thousand kernel weight (TKW) was determined using an optical seed counter (Contador, Pfeuffer GmbH, Germany), with results expressed on a dry-weight basis. Screenings (%) were quantified by passing grain over a standard 2.8-mm slotted sieve and calculating the proportion of grain retained versus discarded. Grain protein content, test weight, and moisture content were measured using a near-infrared spectroscopy instrument (FOSS analytical system) following manufacturer guidelines.

### Data Analyses

#### Stomatal Anatomy & Deep Learning Model

In all seasons, stomatal traits were quantified using a deep-learning-based image analysis pipeline, as described by Chaplin et al., (2025). In brief, 121 images were initially annotated via instance segmentation using Roboflow with the images split into train (60%), validation (30%) and test (10%). Each image was then split into 1280×1280 tiles and exported as both an original and augmented image. Image augmentations included random rotations, flips, contrast and saturation changes. Tiling and augmentation resulted in 822 train images, 420 validation images and 78 test images. YOLOv8 models of varying sizes were trained and evaluated. YOLOv8-M which ran for 119 epochs was selected for further analyses based on performance metrics including precision, recall, and mean average precision (mAP) performance (Table 1). Stomatal traits including SD and guard cell dimensions were extracted using a custom Python script employing Ultralytics and OpenCV. This performed automated stomatal detection, and a fitted ellipse around each stomata enabled anatomical measurements to be estimated (Figures S2 and S3). Guard cell length (GCL) was defined as the major axis of the fitted ellipse, representing the longitudinal dimension of the stomatal pore complex. Guard cell width (GCW) was defined as the minor axis of the fitted ellipse, representing the transverse dimension. Stomatal area (SA) was calculated based on the pixel area of detected stomata, providing an integrative anatomical metric and *g_smax_* was calculated according to Franks and Beerling (2009). To refine *g_smax_* calculations, the conventional assumption that pore length is half guard cell length was tested (Franks and Beerling, 2009; Franks and Farquhar, 2007). The empirically determined value (0.543) was incorporated into *g_smax_* calculations. Full methodological details for stomatal anatomy image analysis are provided in Chaplin et al., (2025) and are available via the GitHub repository: https://github.com/williamtsalter/FieldDinoMicroscopy.

### Phenotypic Visual C Statistical Analyses

Phenotypic data visualisation were performed with R (R Core Team, 2021) using packages dplyr (Wickham et al., 2023), ggplot2 (Wickham, 2016), gridExtra (Baptiste, 2017) and tidyr (Wickham et al., 2024). Statistical analyses were performed with R (R Core Team, 2021) using a linear mixed-effects model (LMM) to account for potential variation among replicates. The model was fitted using the lme4 package in R, with Treatment, Variety, and Surface as fixed effects, and Rep and Leaf as random effects:

*Response Var (e.g.,g_s_) ∼ Treatment x Variety x Surface + (1|Rep) +x (1|Leap)*

Analysis of variance (ANOVA) was performed to assess the significance of fixed effects and their interactions, and post-hoc pairwise comparisons were conducted using the emmeans package, with Tukey’s adjustment for multiple comparisons between separate treatment groups. Packages lme4 (Bates et al., 2015), lmerTest (Kuznetsova et al., 2017), ggplot2 (Wickham, 2016), emmeans (Lenth et al., 2020) and MASS (Venables and Ripley, 2002) were used for analyses.

Prior to model fitting, assumptions of normality and homoscedasticity of residuals were assessed visually. Where assumptions were not met, the Box-Cox transformation (Box and Cox, 1964) was applied using the boxcox function from the MASS package. The optimal transformation parameter, *λ*, was identified by maximising the log-likelihood of a fixed-effects linear model across a range of *λ ϵ* [−2,2]. Based on the estimated *λ*, the most appropriate transformation was applied (e.g., log if *λ* ≈ 0, square root if *λ* ≈ 0.5, cube root if *λ* ≈ 0.33) prior to fitting the mixed-effects model (Wilkinson and Rogers, 1973). Interpretation of the TOS effect should be made with caution, as it may be partially attributable to spatial variation resulting from time of sowing being spatially confounded with block.

### Genotypic Visual & Statistical Analyses

The materials were genotyped using the Illumina Infinium Wheat Barley 40K SNP array (Keeble-Gagnère et al., 2021). The array contains 25,363 wheat-specific and 14,261 barley-specific SNP markers and enables accurate imputation of SNP data. It can therefore be used to infer genotypes at untyped chromosomal locations. for downstream quantitative trait loci (QTL) analysis.

Prior to the QTL analysis, broad-sense heritability was estimated for each trait, trial and surface from the phenotypic model using the approach by Cullis et al. (2006) via the heritable package (Kar and Tanaka, 2025). More specifically, an appropriately transformed phenotypic trait was modelled using experimental design factors (Row, Range, Plot and Sampling Date) as random effects assuming independent Normal distribution with mean zero and constant variance using the asreml R package (Butler et al., 2023). If the estimated heritability was greater than 0.05, then putative QTLs were searched using a whole genome approach using the wgaim package (Taylor and Verbyla, 2011). This approach involves iteratively fitting each marker as fixed effects in a forward selection procedure to select significant markers, while accounting for background level of genetic variation through random additive marker effects using the genomic relationship matrix (Verbyla et al., 2007). The marker was considered significant using the default family-wise error rate of 0.05 and using an exclusion window of 10 mega base pair (Mbp).

## Results

### Season 1 – 200 Genotypes

#### Stomatal Conductance

In season 1, *g*ₛ was significantly affected by sowing time (TOS), leaf surface, and their interaction (p < 0.001) (Table 2). Plants sown earlier (TOS 1) had higher *g*ₛ than those sown later (TOS 2) for both surfaces (abaxial p < 0.01; adaxial p < 0.001), with the highest values on the adaxial surface at TOS 1 (Figure 1A). On average, *g*ₛ declined by 28.9% under delayed sowing (Table S3). Leaf surface had the strongest effect (p < 0.001), with adaxial *g*ₛ consistently exceeding abaxial *g*ₛ (p < 0.001). A significant TOS × surface interaction (p < 0.001) indicated that sowing time influenced surfaces differently; abaxial *g*ₛ declined by 14.8% from TOS1 to TOS2, while adaxial *g*ₛ dropped by 33.9% (Table S3), making the adaxial surface the main contributor to overall gas exchange and the primary driver of reductions under warmer TOS 2 conditions. No significant genotypic variation was detected among the 200 genotypes.

**Figure 1:**
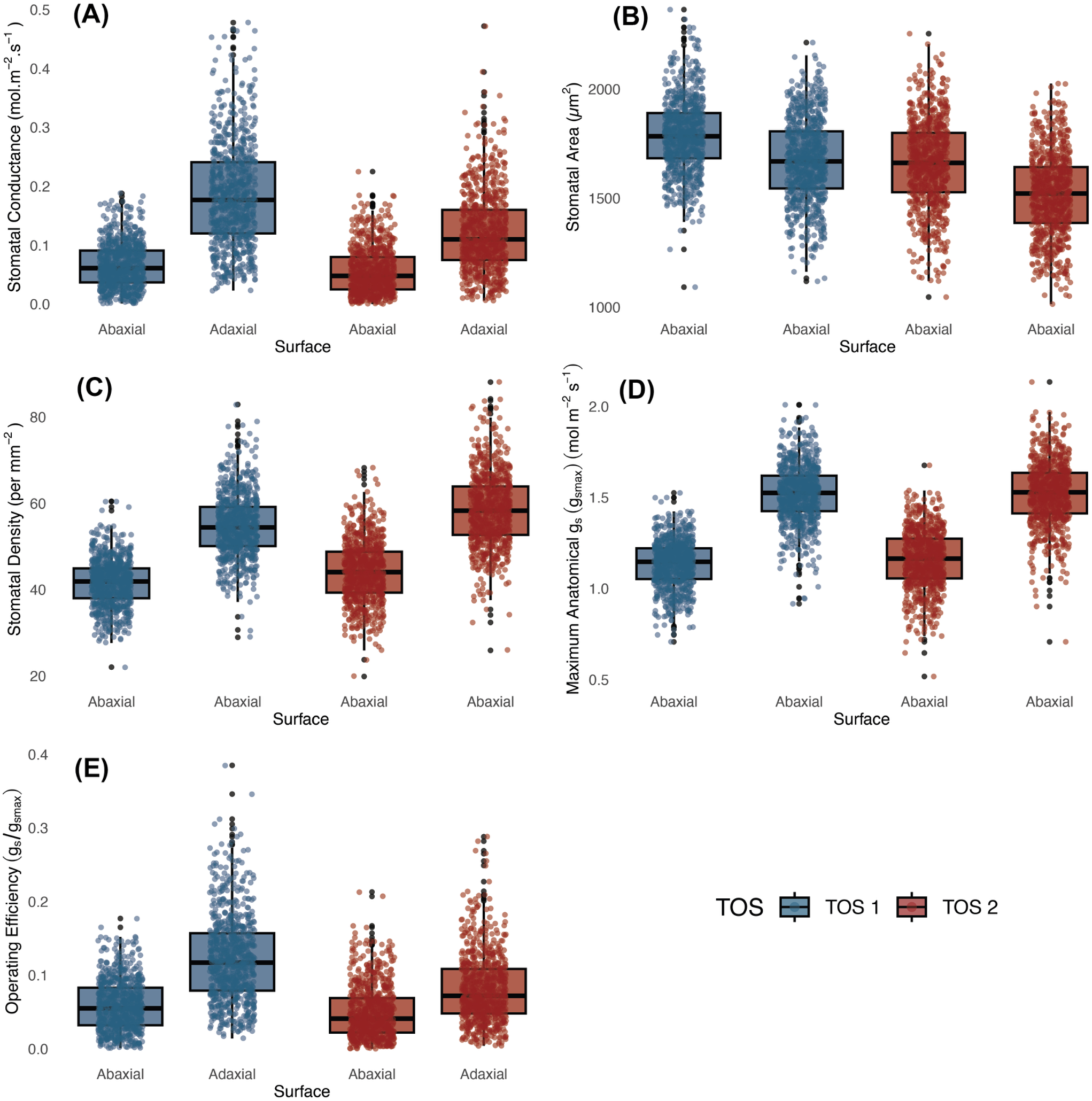
Operational stomatal conductance (*g*_sop_) and anatomical traits across 200 wheat genotypes at two sowing times in season 1. (a) *g*_sop_; (b) stomatal area; (c) stomatal density; (d) maximal anatomical stomatal conductance; and (e) stomatal conductance operating efficiency (*g_se_*). Boxes represent the interquartile range (25th–75th percentiles), with horizontal lines indicating the median. Whiskers denote the minimum and maximum values, and points represent individual observations.

#### Stomatal Anatomy

Guard cell length (GCL) was significantly larger at TOS 1 than TOS 2 (p < 0.001) (Table 2), with strong genotype effects (p < 0.001). A significant TOS × surface interaction (p < 0.01) indicated that sowing time influenced GCL differently across leaf surfaces. Post-hoc tests showed adaxial GCL exceeded abaxial only at TOS 1 (p < 0.001). Mean GCL declined by 5.0% from TOS 1 to TOS 2 (Table S3).

Guard cell width (GCW) was also significantly larger at TOS 1 than TOS 2 (p < 0.001), averaging 3.0% lower in TOS 2 (Table S3). Surface had a strong effect (p < 0.001), with abaxial GCW consistently larger than adaxial (p < 0.001). Genotype also influenced GCW (p < 0.001), but no significant interactions among TOS, surface, and genotype were detected (all p > 0.17), indicating independent effects.

Stomatal area (SA) was also significantly larger at TOS 1 than TOS 2 (p < 0.001), declining by 8.2% (Figure 1B; Table S3). Genotype effects were significant (p < 0.001), while surface showed no main effect (p = 0.211) (Table 2). A TOS × surface interaction (p < 0.01) indicated greater reductions on the adaxial surface between TOS 1 and TOS2 compared to the abaxial.

Stomatal density was significantly affected by sowing time (TOS), genotype, and surface (all p < 0.001) (Table 2; Figure 1C; Figure 2). A significant genotype × surface interaction (p < 0.01) indicated that genotypic effects differed between surfaces, whereas TOS × variety (p = 0.717) and TOS × surface (p = 0.174) interactions were not significant, suggesting TOS effects were consistent across genotypes and surfaces. Post-hoc tests confirmed SD was larger at TOS 2 than TOS 1 for both surfaces (p < 0.001), increasing by 6.4% (abaxial) and 6.7% (adaxial) (Table S3). Adaxial SD exceeded abaxial across treatments (p < 0.001). SD correlated positively with *g*_s_ (R² = 0.1383) and negatively with stomatal area (SA) (R² = 0.321) (Figure S4).

**Figure 2:**
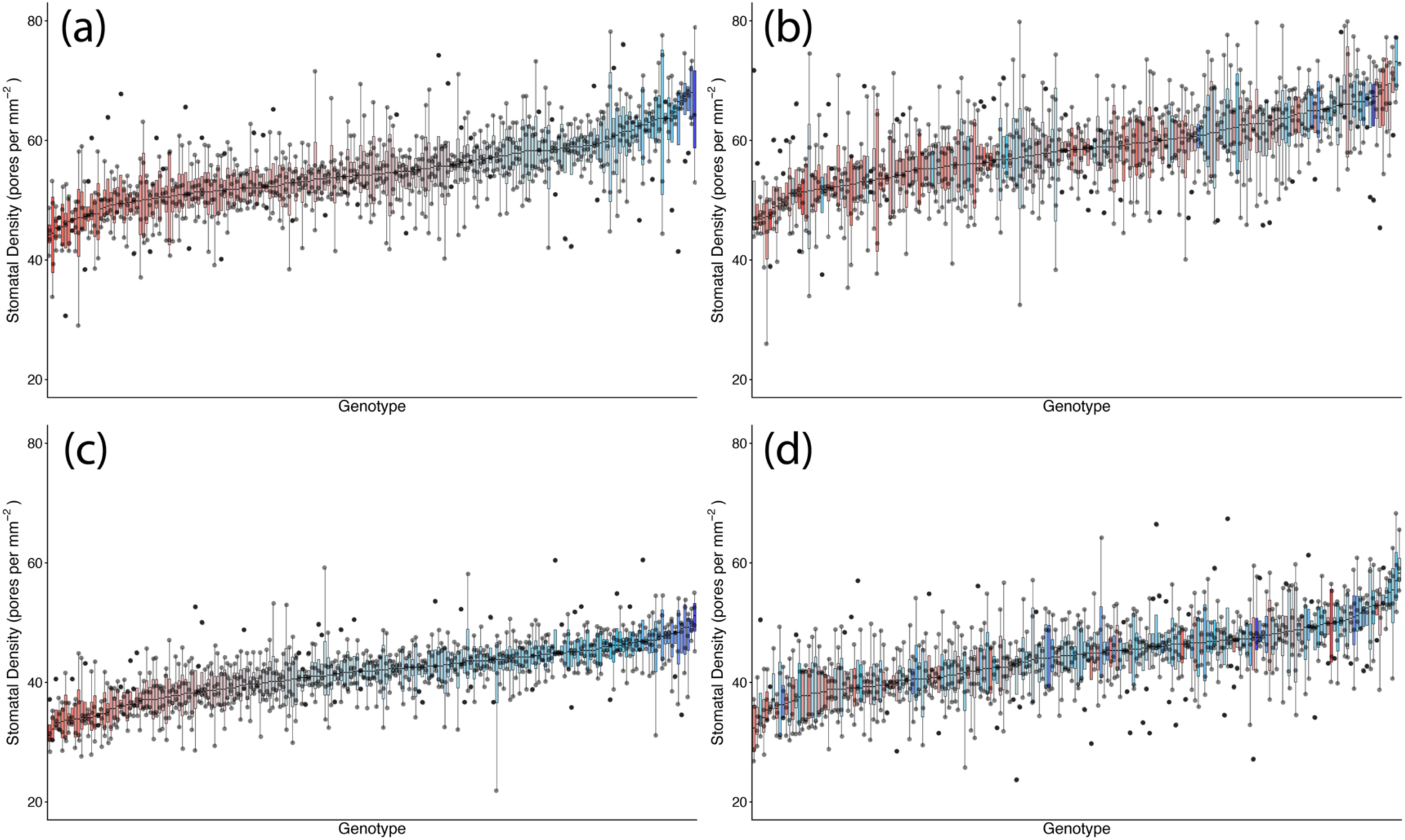
Genotypic distribution of stomatal density across 200 wheat genotypes at two sowing times in season 1. (a) Adaxial surface TOS 1; (b) adaxial surface TOS 2; (c) abaxial surface TOS 1; and (d) abaxial surface TOS 2. Genotypes are ranked by median stomatal density. Colour assigned to each genotype based on TOS 1 stomatal density. Thick horizontal lines within boxes indicate the median and boxes indicate the upper (75%) and lower (25%) quartiles. Whiskers indicate the ranges of the minimum and maximum values. Points indicate individual measurements.

*g*_smax_, estimated from stomatal anatomy, showed no significant effect of sowing time (TOS) but a weak genotypic effect (p < 0.05), indicating differences among genotypes. Median adaxial *g*_smax_ ranged from 1.27–1.84 mol m^-2^ s^-1^ (TOS 1) and 1.18–1.91 mol m^-2^ s^-1^ (TOS 2); abaxial values ranged from 0.85–1.36 mol m^-2^ s^-1^ and 0.84–1.44 mol m^-2^ s^-1^, respectively (Table 2; Figure 3). Surface had a strong effect (p < 0.001) on *g*_smax_, with adaxial estimates consistently higher than abaxial across TOS (p < 0.001) (Figure 1D). A significant variety × surface interaction (p < 0.05) indicated that genotype effects on *g*_smax_ differed between leaf surfaces. No three-way interaction was detected (p = 0.310). *g*_smax_ was positively correlated with measured *g*_s_ (R² = 0.213).

**Figure 3:**
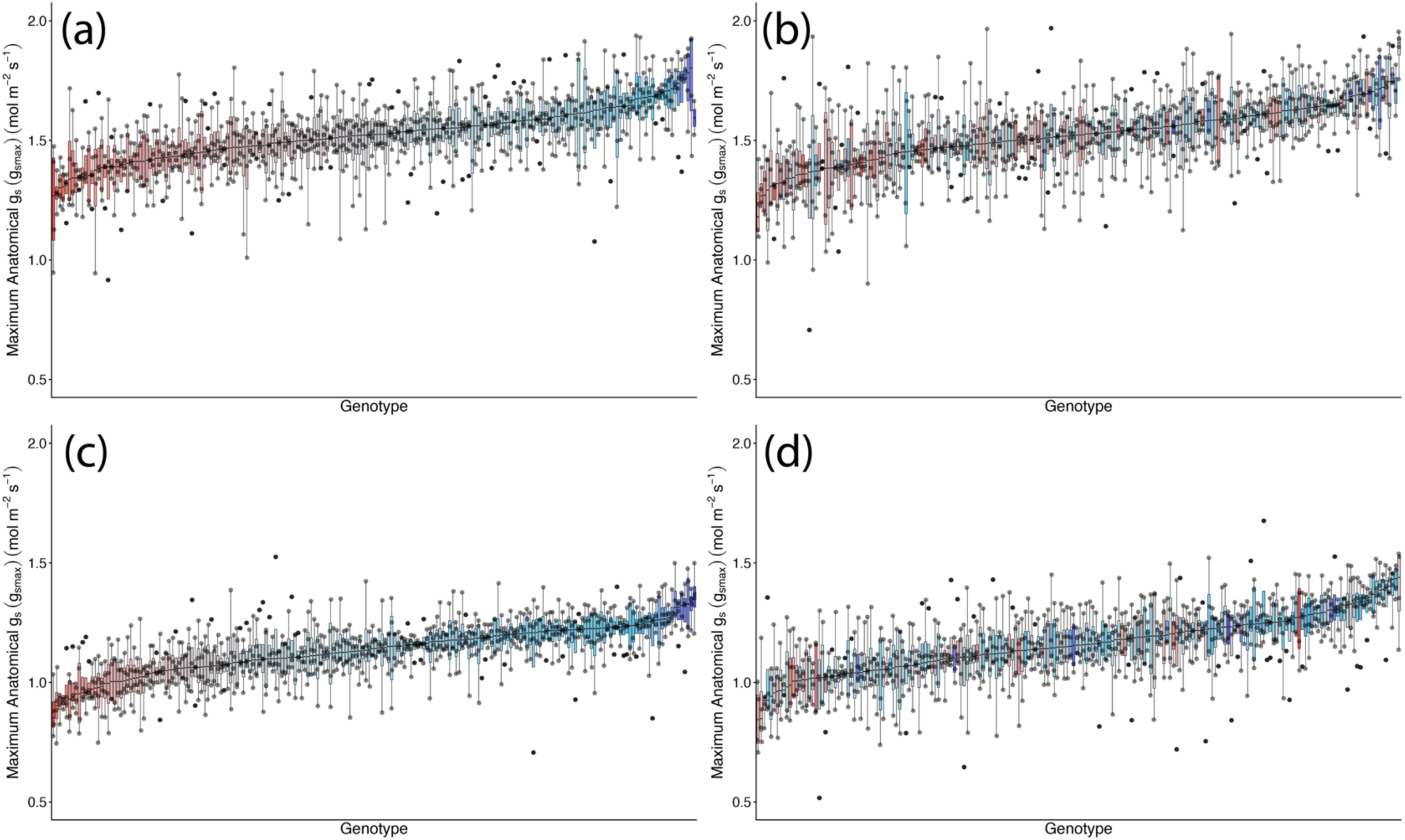
Genotypic distribution of maximum anatomical stomatal conductance, *g_smax_*, across 200 wheat genotypes at two sowing times in season 1. (a) Adaxial surface TOS 1; (b) adaxial surface TOS 2; (c) abaxial surface TOS 1; and (d) abaxial surface TOS 2. Genotypes are ranked by median *g_smax_*. Colour assigned to each genotype based on TOS 1 *g_smax_*. Thick horizontal lines within boxes indicate the median and boxes indicate the upper (75%) and lower (25%) quartiles. Whiskers indicate the ranges of the minimum and maximum values. Points indicate individual measurements.

#### Integrated Stomatal Conductance C Anatomy

When *g*_s_ and *g*_smax_ were combined to calculate *g*_se_ (unitless), significant effects of sowing time (TOS) and surface were detected (p < 0.001) (Table 2; Figure 1E), along with a strong TOS × surface interaction (p < 0.001), indicating that TOS effects varied by leaf surface. Post-hoc tests revealed *g*_se_ was higher at TOS 1 than TOS 2 for both surfaces (p < 0.001), declining by 15.2% on the abaxial and 33.3% on the adaxial surface (Table S3). Across both sowing times, adaxial *g*_se_ exceeded abaxial (p < 0.001). There was no significant variation across genotypes.

#### Yield Parameters

##### Grain Yield

Grain yield was significantly higher at TOS 1 compared with at TOS 2 (p<0.001) (Table 2). Mean grain yield at TOS 1 was 4.62 t/ha while the mean grain yield at TOS 2 was 2.79 t/ha, equating to a 39.6% reduction between TOS. Genotypes also varied significantly with regards to grain yield (p<0.001). Additionally, the TOS x Genotype interaction was significant (p<0.001) suggesting that the effect of TOS on yield varies dependent on genotype. No significant relationships were found between yield and stomatal traits.

#### Genome-Phenome Analyses

Stomatal anatomical traits (GCL, GCW, SA and SD) exhibited heritability estimates of 0.375 to 0.677, while the heritability estimates for stomatal conductance traits (*g*ₛ and *g_se_*) were generally lower, ranging from 0.094 to 0.392, although *g_smax_* was higher ranging from 0.440 to 0.653 (Table 3). While there were no considerable differences in estimated heritability across TOS for abaxial surfaces, all heritability estimates were lower at TOS 2 for the adaxial surface.

**Table 3:**
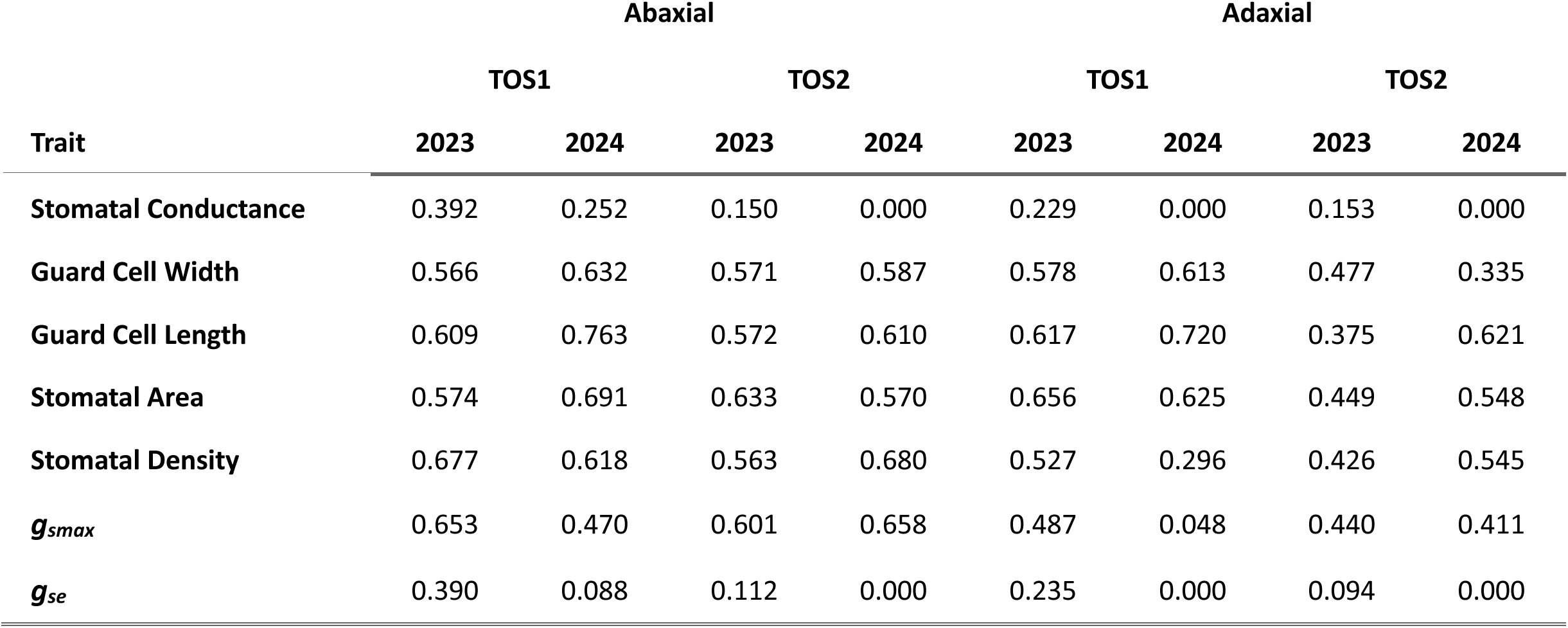
Broad-sense heritability estimates for each trait, arranged by surface, TOS and year.

At least one putative QTL candidate was identified for each trait for every surface and TOS, except *g_smax_* for the abaxial surface at TOS 2 and *g_s_* and *g_se_* for the adaxial surface (Table 4). In total, 96 putative QTLs (49 abaxial and 47 adaxial) were found across stomatal anatomical traits while 10 were identified for stomatal conductance traits (6 for *g_s_* and 4 for *g_se_*). All QTL candidates for *g_s_* and *g_se_* were found for the abaxial surface, and the majority were at TOS 2. All traits appear to be explained by several chromosomal locations across the whole genome (Figure 4; Figure S5; Table S4). Two markers have noticeably larger LOD scores than others: one at chromosome 6B for GCW (LOD = 13.8) and one at chromosome 5A for GCL (LOD = 12.3) (Figure S5). Chromosome 7B appears to contain several closely positioned QTLs for stomatal anatomical traits (see full list of putative QTLs in Table S5).

**Figure 4:**
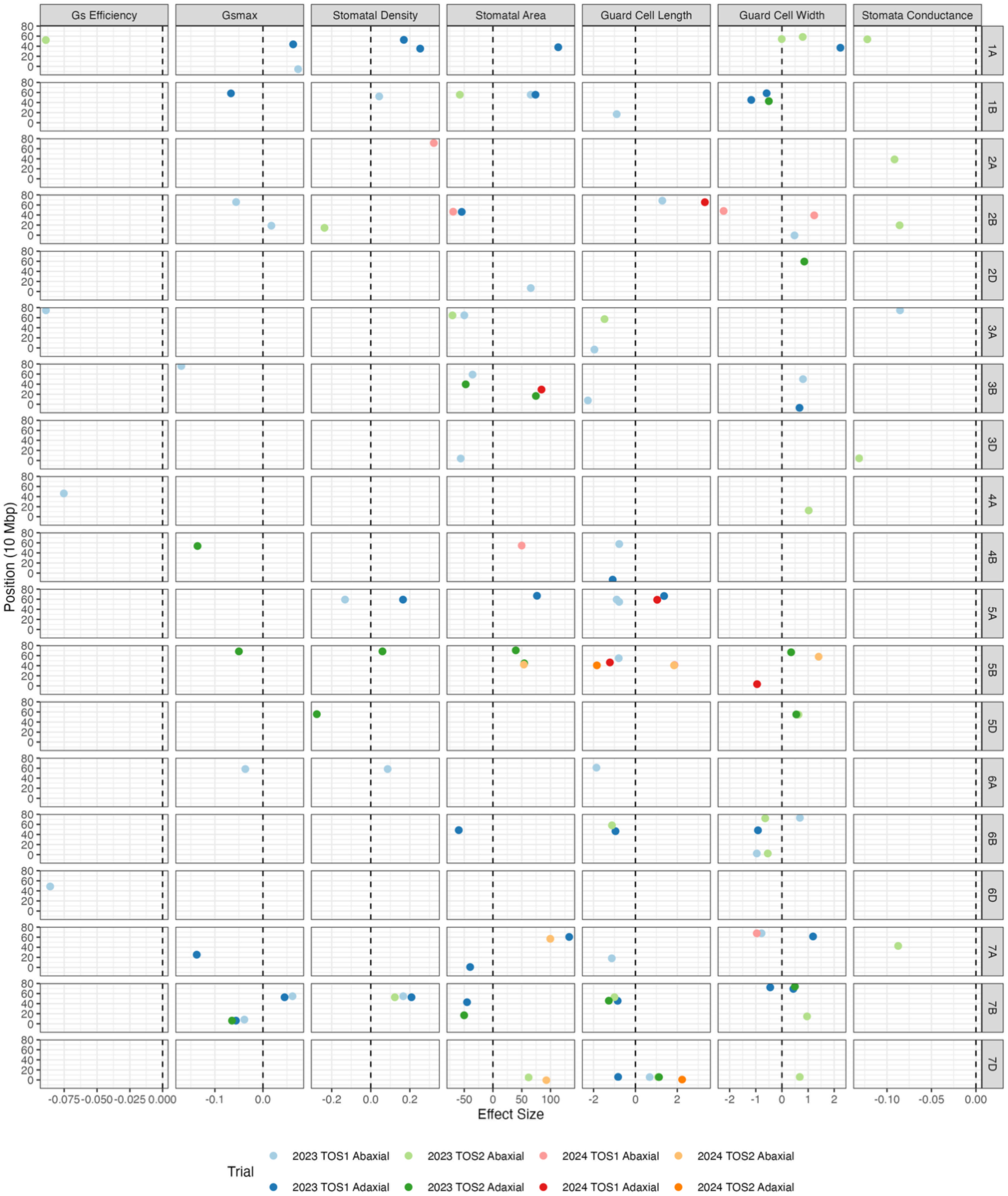
Effective sizes of putative QTLs by trait and chromosome. The scatterplot shows the effect sizes of putatiative QTLs identified by chromosome and position (in mega base pairs), with each color corresponding to a specific year, TOS and surface. The vertical dashed lines are the y-intercepts. The points are slightly jittered so the points are visible.

**Table 4:** The number of putative QTL candidates by year, TOS, and surface for each trait. The “.” represents 0.

| Trait | Abaxial |  |  |  | Adaxial |  |  |  |
| --- | --- | --- | --- | --- | --- | --- | --- | --- |
|  | TOS1 |  | TOS2 |  | TOS1 |  | TOS2 |  |
|  | 2023 | 2024 | 2023 | 2024 | 2023 | 2024 | 2023 | 2024 |
| Stomatal Conductance | 1 | . | 5 | . | . | . | . | . |
| Guard Cell Width | 5 | 3 | 8 | 1 | 8 | 1 | 5 | . |
| Guard Cell Length | 11 | 1 | 4 | 1 | 5 | 3 | 2 | 2 |
| Stomatal Area | 5 | 2 | 3 | 3 | 8 | 1 | 5 | . |
| Stomatal Density | 4 | 1 | 2 | . | 4 | . | 2 | . |
| $g_{smax}$ | 7 | . | . | . | 5 | . | 3 | . |
| $g_{se}$ | 3 | . | 1 | . | . | . | . | . |

### Season 2 – 50 Genotypes

#### Stomatal Conductance

In season 2, *g*ₛ was significantly affected by genotype and leaf surface (both p < 0.001), along with several interactions (Table 5). However, the linear mixed-effects model showed no main effect of sowing time (TOS) (p = 0.325) (Figure 5A). Leaf surface exerted the strongest influence on *g*ₛ (p < 0.001), with adaxial *g*ₛ consistently higher than abaxial *g*ₛ (p < 0.001).

**Figure 5:**
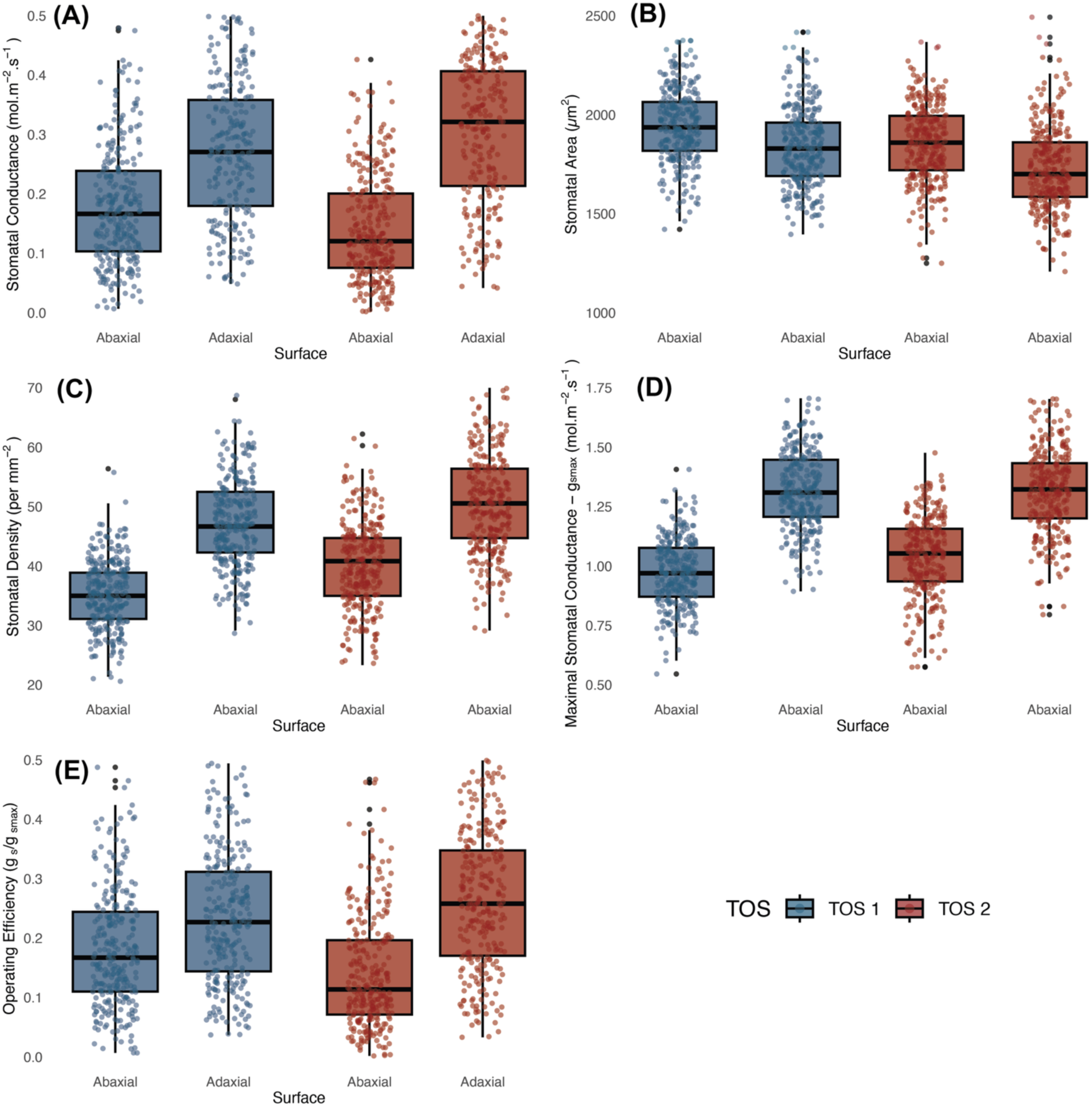
Operational stomatal conductance (*g*_sop_) and anatomical traits across 200 wheat genotypes at two sowing times in season 2. (a) *g*_sop_; (b) stomatal area; (c) stomatal density; (d) maximal anatomical stomatal conductance; and (e) stomatal conductance operating efficiency (*g_se_*). Boxes represent the interquartile range (25th–75th percentiles), with horizontal lines indicating the median. Whiskers denote the minimum and maximum values, and points represent individual observations.

**Table 5:**
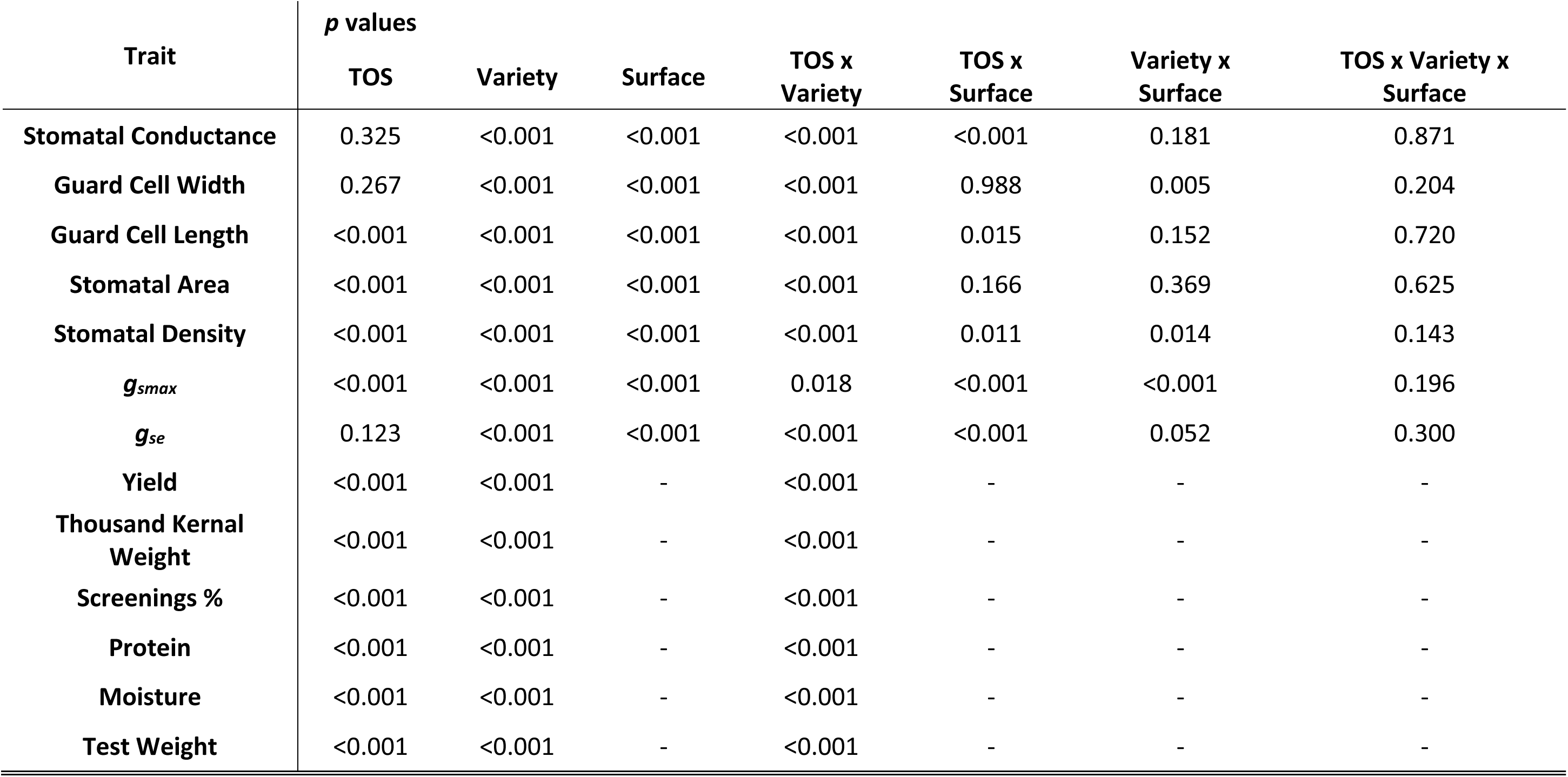
Linear mixed model ANOVA p values for TOS, Variety and Surface on traits of wheat from 2024 field data.

| Trait | *p* values |  |  |  |  |  |  |
| --- | --- | --- | --- | --- | --- | --- | --- |
|  | TOS | Variety | Surface | TOS x Variety | TOS x Surface | Variety x Surface | TOS x Variety x Surface |
| **Stomatal Conductance** | 0.325 | <0.001 | <0.001 | <0.001 | <0.001 | 0.181 | 0.871 |
| **Guard Cell Width** | 0.267 | <0.001 | <0.001 | <0.001 | 0.988 | 0.005 | 0.204 |
| **Guard Cell Length** | <0.001 | <0.001 | <0.001 | <0.001 | 0.015 | 0.152 | 0.720 |
| **Stomatal Area** | <0.001 | <0.001 | <0.001 | <0.001 | 0.166 | 0.369 | 0.625 |
| **Stomatal Density** | <0.001 | <0.001 | <0.001 | <0.001 | 0.011 | 0.014 | 0.143 |
| ***g<sub>smax</sub>*** | <0.001 | <0.001 | <0.001 | 0.018 | <0.001 | <0.001 | 0.196 |
| ***g<sub>se</sub>*** | 0.123 | <0.001 | <0.001 | <0.001 | <0.001 | 0.052 | 0.300 |
| **Yield** | <0.001 | <0.001 | - | <0.001 | - | - | - |
| **Thousand Kernal Weight** | <0.001 | <0.001 | - | <0.001 | - | - | - |
| **Screenings %** | <0.001 | <0.001 | - | <0.001 | - | - | - |
| **Protein** | <0.001 | <0.001 | - | <0.001 | - | - | - |
| **Moisture** | <0.001 | <0.001 | - | <0.001 | - | - | - |
| **Test Weight** | <0.001 | <0.001 | - | <0.001 | - | - | - |

Genotypic variation was substantial (p < 0.001) among the 50 wheat genotypes (Figure 6). Median ranges with median adaxial *g*ₛ ranging from 0.130–0.605 mmol m^-2^ s^-1^ (TOS 1) and 0.162–0.620 mmol m^-2^ s^-1^ (TOS 2) and abaxial *g*ₛ from 0.072–0.291 mmol m^-2^ s^-1^ (TOS 1) and 0.044–0.270 mmol m^-2^ s^-1^ (TOS 2).

**Figure 6:**
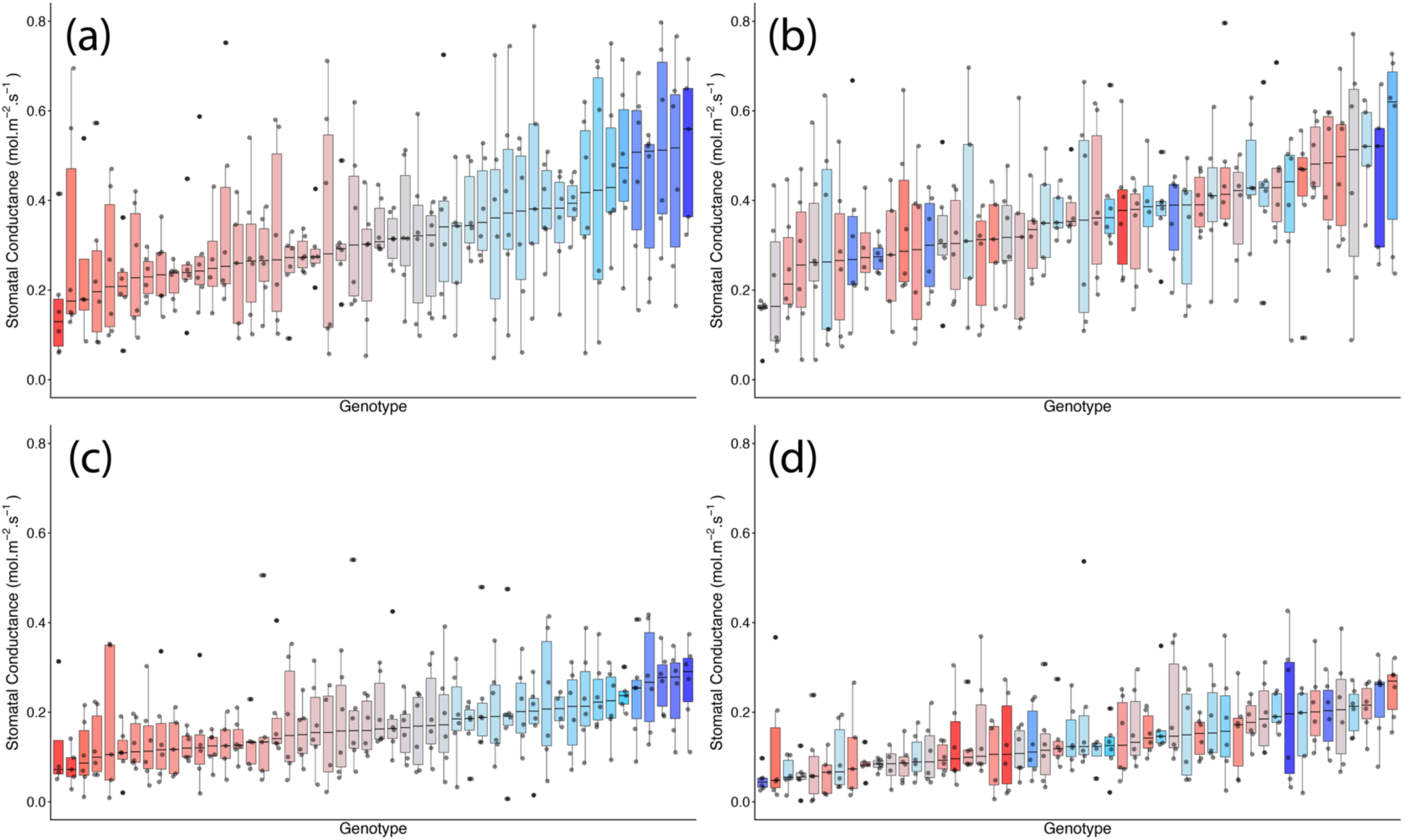
Genotypic distribution of stomatal conductance (*g_s_*) across 50 wheat genotypes at two sowing times in season 2. (A) Adaxial surface TOS 1; (B) adaxial surface TOS 2; (C) abaxial surface TOS 1; and (D) abaxial surface TOS 2. Genotypes are ranked by median *g_s_*. Colour assigned to each genotype based on TOS 1 *g_s_*. Thick horizontal lines within boxes indicate the median and boxes indicate the upper (75%) and lower (25%) quartiles. Whiskers indicate the ranges of the minimum and maximum values. Points indicate individual measurements.

Although TOS had no overall effect, a significant TOS × genotype interaction (p < 0.001) indicated non-uniform genotypic responses, with some lines showing greater declines under later sowing. Similarly, a significant TOS × surface interaction (p < 0.001) revealed that sowing time influenced surfaces differently, with post-hoc tests confirming a stronger TOS effect on adaxial *g*ₛ.

#### Stomatal Anatomy

GCL varied significantly among genotypes (p < 0.001), between sowing times (p < 0.001), and across leaf surfaces (p < 0.001) (Table 5). Post-hoc tests showed GCL was higher at TOS 1 than TOS 2 for both surfaces (p < 0.001). At TOS 1, adaxial GCL exceeded abaxial (p < 0.001), while no surface difference was detected at TOS 2 (p = 0.167). Significant TOS × variety (p < 0.001) and TOS × surface (p < 0.05) interactions indicate that sowing time effects varied by genotype and surface.

GCW was significantly affected by leaf surface and genotype (both p < 0.001) but not by sowing time (Table 5). Post-hoc tests confirmed that GCW was consistently higher on the abaxial surface than the adaxial surface at both TOS (p < 0.001). Significant TOS × genotype (p < 0.001) and genotype × surface (p < 0.01) interactions indicate that sowing time effects varied among genotypes and that surface contributed to genotypic differences in GCW.

SA was significantly affected by sowing time and leaf surface (both p < 0.001) (Table 5; Figure 5B), with strong genotypic variation (p < 0.001). SA ranged from 1421–2602 µm^2^ (abaxial) and 1396–2661 µm^2^ (adaxial) at TOS 1, and from 1250–2367 µm^2^ (abaxial) and 1208–2493 µm^2^ (adaxial) at TOS 2. Post-hoc tests confirmed SA was higher on the abaxial surface than the adaxial surface at both sowing times (p < 0.001) and higher at TOS 1 than TOS 2 for both surfaces (p < 0.001), with a 4.7% decline on the abaxial surface and 6.2% on the adaxial surface between TOS1 and TOS 2 (Table S6). A significant TOS × variety interaction (p < 0.001) indicates genotype-dependent responses to sowing time.

SD was significantly affected by genotype, sowing time (TOS), and leaf surface (all p < 0.001) (Table 5). Post-hoc tests confirmed that adaxial SD exceeded abaxial SD at both sowing times (p < 0.001) and that SD was higher at TOS 2 than TOS 1 for both surfaces (p < 0.001). The lowest SD occurred on the abaxial surface at TOS 1, significantly lower than all other groups (p < 0.001) (Figure 5C). On average, SD increased from 35.4 to 40.2 mm^-2^ on the abaxial surface and from 46.9 to 50.4 mm^-2^ on the adaxial surface between TOS 1 and TOS 2, representing a 13.5% and 7.3% increase, respectively (Table S6). Significant TOS × variety (p < 0.001), TOS × surface (p < 0.05), and variety × surface (p < 0.05) interactions indicate that SD variation is both genotype- and surface-dependent (Figure 7). A moderate negative correlation was observed between SD and stomatal area (SA) (R^2^ = 0.312).

**Figure 7:**
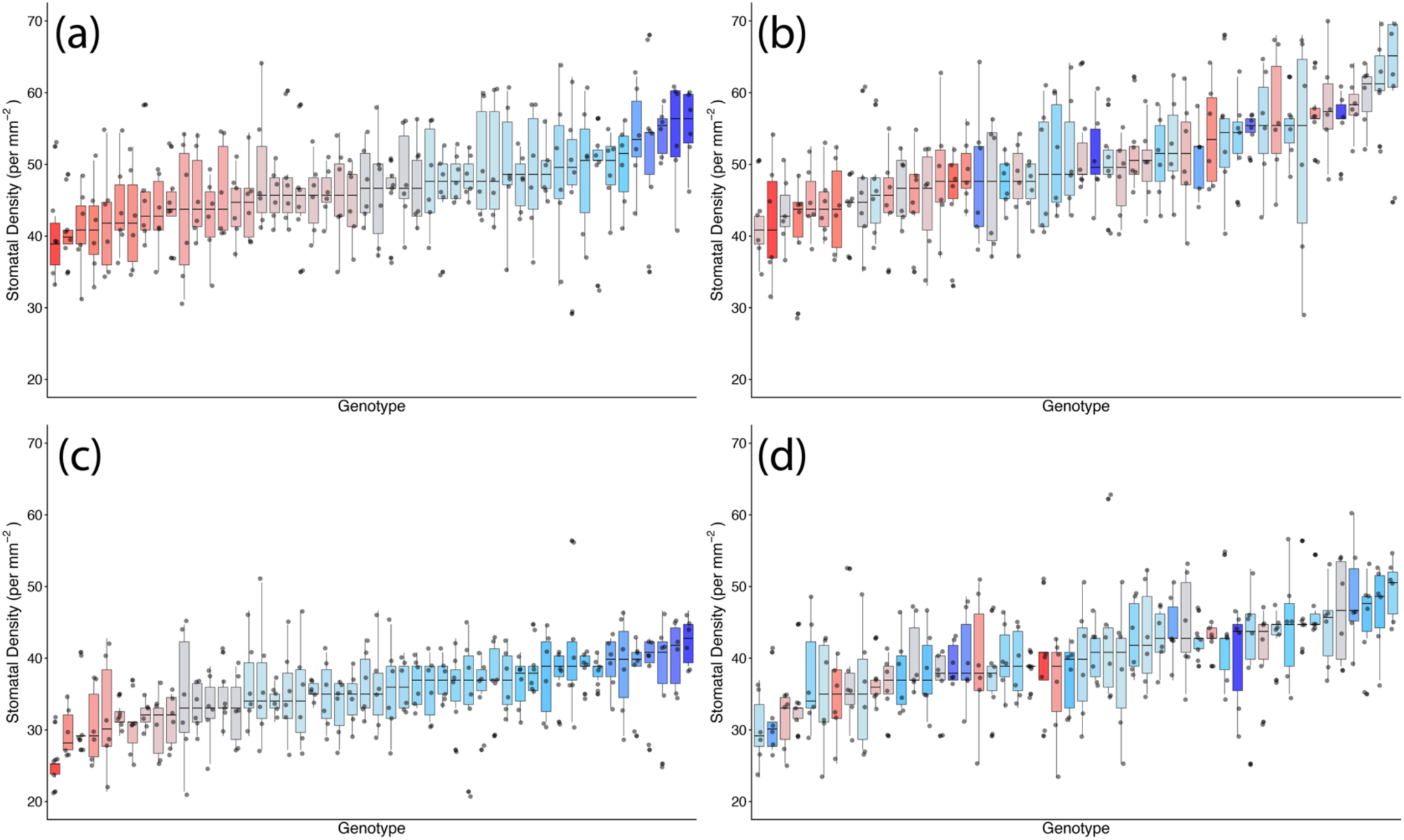
Genotypic distribution of stomatal density across 50 wheat genotypes at two sowing times in season 2. (A) Adaxial surface TOS 1; (B) adaxial surface TOS 2; (C) abaxial surface TOS 1; and (D) abaxial surface TOS 2. Genotypes are ranked by median stomatal density. Colour assigned to each genotype based on TOS 1 stomatal density. Thick horizontal lines within boxes indicate the median and boxes indicate the upper (75%) and lower (25%) quartiles. Whiskers indicate the ranges of the minimum and maximum values. Points indicate individual measurements.

*g*_smax_ varied significantly among genotypes, sowing times (TOS), and leaf surfaces (all p < 0.001) (Table 5; Figure 8). Post-hoc tests showed adaxial *g*_smax_ was consistently higher than abaxial at both sowing times (p < 0.001). While adaxial *g*_smax_ remained unchanged between TOS, abaxial *g*_smax_ increased by 7.1% at TOS 2 compared with TOS 1 (p < 0.001) (Figure 5D). On average, abaxial *g*_smax_ rose from 0.97 to 1.04 mol m^-2^ s^-1^, whereas adaxial *g*_smax_ was stable at 1.32 mol m^-2^ s^-1^ across sowing times. Significant TOS × genotype (p < 0.05), TOS × surface (p < 0.001), and genotype × surface (p < 0.001) interactions indicate that sowing time effects on *g*_smax_ were genotype- and surface-dependent, with leaf surface modulating genotypic responses.

**Figure 8:**
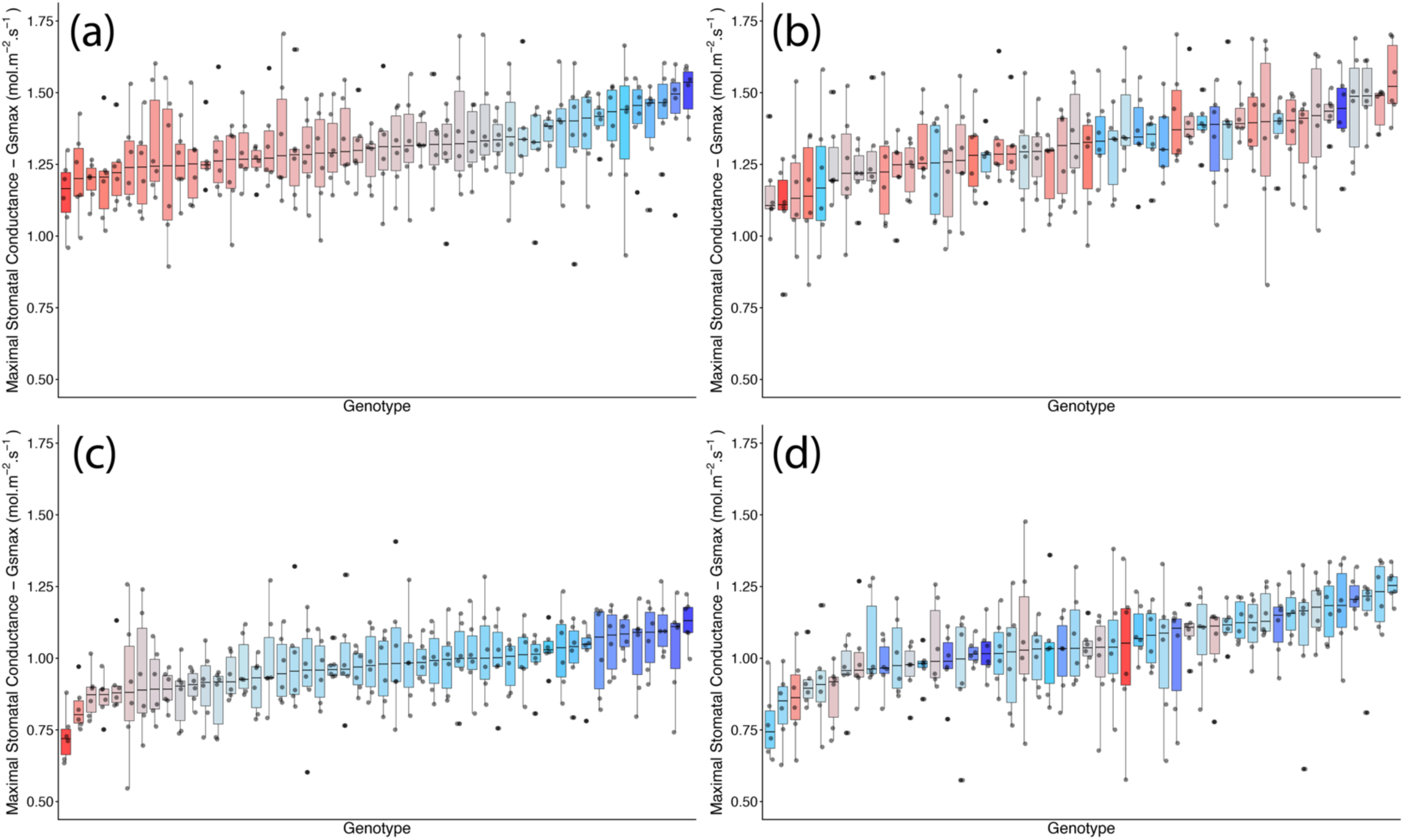
Genotypic distribution of maximum anatomical stomatal conductance, *g_smax_*, across 50 wheat genotypes at two sowing times in season 2. (A) Adaxial surface TOS 1; (B) adaxial surface TOS 2; (C) abaxial surface TOS 1; and (D) abaxial surface TOS 2. Genotypes are ranked by median *g_smax_*. Colour assigned to each genotype based on TOS 1 *g_smax_*. Thick horizontal lines within boxes indicate the median and boxes indicate the upper (75%) and lower (25%) quartiles. Whiskers indicate the ranges of the minimum and maximum values. Points indicate individual measurements.

#### Integrated Stomatal Conductance C Anatomy

Significant effects of leaf surface and genotype on *g*_se_ were detected (both p < 0.001) (Table 5). Post-hoc tests confirmed that adaxial *g*_se_ was consistently higher than abaxial across both sowing times (p < 0.001). No overall decline in *g*_se_ was observed under heat stress (Figure 5E). On average, *g*_se_ decreased slightly on the abaxial surface from 0.18 at TOS 1 to 0.14 at TOS 2, while adaxial *g*_se_ increased from 0.26 to 0.29 across sowing times. Significant TOS × genotype (p < 0.001) and TOS × surface (p < 0.05) interactions indicate that sowing time effects on *g*_se_ were modulated by both genotype and leaf surface.

#### Yield Parameters

##### Grain Yield

Later sowing time significantly reduced yield (p < 0.001). Mean grain yield declined from 5.88 t ha^-1^ at TOS 1 to 3.48 t ha^-1^ at TOS 2, representing a 40.8% reduction (Table S6). Yield also varied significantly among the 50 genotypes (p < 0.001) (Table 5), and a significant TOS × genotype interaction (p < 0.001) indicates that the impact of sowing time was genotype-dependent. No significant relationships were detected between yield and stomatal traits in season 2.

##### Thousand Kernel Weight

Thousand kernel weight (TKW) was significantly lower at TOS 2 than at TOS 1 (p<0.001). There was also significant variation in TKW across the 50 genotypes (p<0.001) (Table 5). Additionally, the TOS x genotype interaction was significant (p<0.001) suggesting that the effect of TOS on TKW varies dependent on genotype.

##### Screenings Percentage

TOS significantly affected screenings percentage (p<0.001), 88.4% higher at TOS 2 than at TOS 1 (p<0.001) (Table S6). There was also significant variation in screenings percentage across the 50 genotypes (p<0.001) (Table 5). Additionally, the TOS x genotype interaction was significant (p<0.001) suggesting that the effect of TOS on screenings percentage varies dependent on genotype.

##### Protein, Moisture & Test Weight

Protein, moisture and test weight were all significantly affected by TOS (p<0.001) and genotype (p<0.001) (Table 5).

#### Genome-Phenome Analyses

Similar to S1, stomatal anatomical traits (GCL, GCW, SA and SD) had the highest heritability estimates (0.296 to 0.763). *g_smax_* had an estimated heritability ranging from 0.048 to 0.658. Stomatal conductance traits, *g*ₛ and *g_se_*, had lower heritability estimates, ranging from 0.089 to 0.252, amongst trials that included detectable heritability estimates (Table 3). Noticeably, heritability estimates for *g*ₛ and *g_se_* were only detectable at TOS1 for the abaxial surface.

A total of 19 putative QTLs (12 for abaxial and 7 for adaxial) were found across stomatal anatomical traits, while none were found for stomatal conductance traits (Table 4). Chromosome 5B contains several potential QTLs for GCL across both TOS 1 and TOS 2 for either surface (Figure 4; Figure S5; Table S4). A marker in chromosome 2B had the largest LOD score (18.5) for GCW (Figure S5). In addition, one putative QTL is also found in chromosome 5B for GCW for each surface. Six QTL candidates from S1 were detected in similar chromosomal regions in S2: stable QTL were reaffirmed on chromosomes 2B for SA, 2B and 5A for GCL, and 7A for GCW, while co-localised QTL across seasons were identified on chromosomes 5B (SA in S1 and GCL in S2) and 4B (*g_smax_* in S1 and SA in S2). The full list of QTL candidates is listed in Table S5.

## Discussion

This study reveals the complex interplay between environmental conditions, stomatal anatomy, physiological responses and their genetic underpinning across a diverse wheat panel, comprising 200 genotypes. We provide robust evidence on how *g_s_*, stomatal anatomy, operating efficiency, and ultimately, grain yield are influenced by sowing time and genotype. The findings underscore the critical role of stomatal traits in mediating plant adaptation to higher temperatures and associated abiotic stress. Identifying putative QTL that are consistent across seasons and associated with multiple traits, particularly when clustered on a single chromosome (7B), highlights their potential as valuable breeding targets.

### Stomatal Physiology

In S1, *g*_s_ was strongly influenced by TOS and leaf surface, alongside significant TOS x Variety and TOS x Surface interactions. Earlier sowing (TOS 1) conferred higher *g*_s_ values compared to later sowing (TOS 2) leading to a decline of 28.9%, likely reflecting more favourable cooler conditions. This is consistent with previous findings highlighting the sensitivity of *g_s_* to thermal stress, particularly in cereals, where later sowing compresses reproductive phases into hotter conditions, limiting carbon assimilation and translocation potential (Mahdavi et al., 2021; Ramya et al., 2016; Schoppach et al., 2020).

This trend was not observed in S2, with *g_s_* not significantly different between sowing times. The divergence in responses between S1 and S2 likely reflects a combination of inter-annual environmental variation (e.g., heat intensity) and genotypic plasticity. In S1, maximum temperatures in TOS 2 rose to an average of 27.2°C compared with an average maximum of 25.1°C at TOS 2 in S2. The less severe heat stress in S2 likely buffered plants against significant declines in *g_s_* under later sowing. Additionally, the lack of a TOS effect on *g*_s_ in S2 may relate to the inherent stress tolerance of the germplasm used. Previous studies have shown that certain heat-tolerant wheat cultivars can maintain or even increase *g*_s_ under high temperatures as part of their adaptive strategy to sustain photosynthesis under stress (Abdelhakim et al., 2021; Mirosavljević et al., 2021; Ramya et al., 2021). Genotype-specific resilience may have reduced population-level differences between sowing times in S2, highlighting the importance of understanding trait plasticity and genotype-by-environment interactions in breeding programs. *g_s_* responses are not only genotype-dependent but are highly context-specific, with environmental factors such as heat intensity and timing influencing *g_s_* responses to stress.

The adaxial leaf surface exhibited consistently higher *g_s_* than the abaxial surface in both years, reinforcing its role as the primary contributor to leaf gas exchange (Pinto et al., 2025; Shahinnia et al., 2016; Wall et al., 2023, 2022). Similarly, the TOS × Surface interaction suggests that genotypes differ in how stomatal behaviour is partitioned between surfaces under environmental stress, with some showing more balanced or adaptive surface-specific responses. Additionally, the effect of TOS on *g_s_* was most substantial on the adaxial surface in both years, highlighting the functional responsiveness of the adaxial surface in driving leaf gas exchange. This aligns with evidence that adaxial stomata – often more responsive, functionally active and present at higher densities – play a more critical role in regulating photosynthesis and transpiration than their abaxial counterparts, particularly under higher temperatures (Pinto et al., 2025; Wall et al., 2022).

### QTL for Stomatal Physiology

The significant TOS × Variety interaction observed across both datasets indicates that genotypes differ not only in their baseline levels of *g_s_* but also in their plasticity to adjust *g_s_* in response to higher temperatures. Such genotype-dependent plasticity demonstrates that genetic control of *g_s_* is environment-specific rather than constitutive, reflecting multiple physiological strategies for heat tolerance. QTL for *g_s_* have previously been identified in wheat under abiotic stress. Wang et al., (2015) identified QTL for *g_s_* under abiotic stress conditions, including a locus on chromosome 2B, which have confirmed in our study, with a particularly high LOD score (18.53). Wang et al., (2015) additionally reported a QTL for *g_s_* on chromosome 7B, a region which in our study was associated primarily with anatomical traits rather than conductance. Notably, chromosome 7B harboured a high density of clustered QTL for multiple stomatal anatomical traits in our analysis, suggesting that this region likely influences *g_s_* indirectly via stomatal anatomy, with the functional expression of these traits potentially contingent on environmental context or developmental timing. The higher heritability estimates for *g_s_* on the abaxial surface, particularly at TOS 1, further indicate that genetic control of stomatal function is surface-specific and may be more stable under less extreme stress. Notably, all six identified QTL candidates for *g_s_* were associated with the abaxial surface and distributed across multiple chromosomes, reinforcing the polygenic nature of stomatal regulation (Figure 4). However, as these QTL were derived from a single trial, their consistency across environments remains to be validated before deployment in breeding programs.

These findings suggest that both intrinsic stomatal capacity and dynamic regulation differ between genotypes and are modulated by environmental context, particularly sowing time. This is an important opportunity for breeding programs; beyond targeting absolute *g_s_* values, selection should focus on genotypes that maintain favourable *g_s_* profiles across sowing times, those that show optimal adjustment in specific conditions or those with the most advantageous surface-level allocation and subsequent stomatal responsiveness. Integrating these traits into selection frameworks could improve both productivity and resilience in wheat, particularly in the face of increasing climate variability.

### Stomatal Anatomy

Anatomical traits such as GCL, SA, and SD were significantly influenced by sowing time and genotype in both seasons, with consistent and strong effects of leaf surface, particularly for SD and GCW. The adaxial surface exhibited significantly higher SD than the abaxial surface in both years, reinforcing established patterns in wheat and other grasses (McAusland et al., 2021; Shahinnia et al., 2016; Wall et al., 2023, 2022). Significant genetic determination of SD and surface-specific stomatal distribution suggests functional specialisation, with the adaxial surface contributing more prominently to gas exchange under favourable conditions (Samantara et al., 2025).

Earlier sowing (TOS 1) was consistently associated with larger guard cells and a greater SA, yet by contrast, later sowing (TOS 2) resulted in smaller stomata with reduced SA, found at higher densities. These anatomical shifts, consistent with prior literature, likely represent adaptive developmental responses to increased heat and evaporative demand later in the season (Pinto et al., 2025). The strongest reductions in SA occurred on the adaxial surface, suggesting that it is also the most plastic and environmentally responsive (Hõrak, 2025; Wall et al., 2022).

A negative correlation between SA and SD was observed across both seasons, consistent with a well-established size–density trade-off (Drake et al., 2013; Haworth et al., 2023; Lawson and Blatt, 2014; Pinto et al., 2025). Smaller stomata at higher densities are often associated with faster stomatal kinetics, improving responsiveness to environmental fluctuations, thus enabling plants to optimise the balance between carbon uptake and water loss on short timescales (Drake et al., 2013; McAusland et al., 2016). This can be especially valuable under later sowing conditions where high VPDs and heat spikes create a dynamic stress landscape as a dense arrangement of small stomata may allow tighter and faster control over transpirational water loss without sacrificing photosynthetic capacity. Such an approach aligns with efforts to enhance both intrinsic WUE and the speed and precision of gas exchange regulation, a dual pathway aimed at improving resilience in climate-challenged cropping systems. The elevated adaxial SD observed in our study likely contributes to the higher *g_s_* and *g_smax_* values seen on this surface across both seasons. This anatomical configuration of smaller stomata at greater density may underpin the adaxial surface’s superior capacity for both baseline gas exchange and for dynamic and rapid adjustment under stress (Faralli et al., 2019; McAusland et al., 2021; Pflüger, 2025). Under stressful conditions, the ability to rapidly close stomata can reduce water loss while equally rapid reopening can ensure photosynthetic recovery once conditions improve.

The benefits of high SD are not universal amongst stress types and, in some cases, lower SD has been associated with enhanced water-use efficiency and drought tolerance while negative relationships between SD and *g_smax_* with grain yield have also been reported in wheat, likely reflecting greater total water loss through more numerous adaxial stomata (Dunn et al., 2019; Li et al., 2023, 2017; Samantara et al., 2025). These varying trends where contrasting traits potentially confer resilience in different scenarios suggest that the optimal anatomical ideotype is likely context-dependent and stress-type specific, requiring functional integration with physiological control mechanisms including efficient stomatal kinetics and regulation to fully realise its benefits and confer resilience in different environments.

A higher adaxial *g_smax_* aligns with prior literature, affirming the adaxial surface as the dominant driver of leaf gas exchange (Samantara et al., 2025). That abaxial *g_smax_* increased under stress at TOS 2 is consistent with Pinto et al., (2025). This suggests an adaptive anatomical shift whereby the abaxial surface, typically contributing less to gas exchange, increases its capacity when adaxial function is constrained by heat stress, potentially enhancing transpirational cooling or buffering against reductions in total foliar carbon assimilation (Wall et al., 2022).

The positive correlation between *g_smax_* and *g_sop_* confirms that anatomical potential influences, but does not fully determine, functional gas exchange outcomes (McAusland et al., 2021). *g_sop_* reflects a dynamic interplay where anatomical capacity sets the upper limit, while short-term environmental conditions and stomatal regulation determine its actual expression. This underscores the importance of integrating both anatomical and physiological traits in breeding strategies. Breeders should consider SA, SD, and *g_smax_* alongside dynamic stomatal control to develop wheat varieties that can efficiently balance carbon assimilation and water use under heat stress. That being said, phenotyping dynamic stomatal responses and kinetics *in-situ* under field conditions at scale remains a significant challenge, currently limiting the incorporation of these dynamic stomatal traits into breeding pipelines.

### QTL for Stomatal Anatomy

QTL for stomatal anatomical traits including stomatal density and size have been previously mapped in wheat, some with pleiotropic loci for yield. Of the 125 putative QTL we found across seven stomatal traits, 42 overlapped with the same chromosomal regions for the same traits previously reported by Liu et al. (2025), Shahinnia et al. (2016) and Wang et al. (2016) (Table S7). However, while overlapping chromosomal regions were identified, it is important to note that due to different genotyping platforms, the specific genomic loci of the markers cannot be mapped across different studies, thus it cannot be determined whether the 42 overlapping regions related to the same precise QTL. Furthermore, a QTL on chromosome 2B for GCW, associated with the marker with the highest LOD score (LOD = 18.5), was also reported by Liu et al. (2025), suggesting chromosome 2B has essential genes involved in the regulation pore complex size.

Stable QTL detected across seasons are strong candidates for marker-assisted selection due to their robustness to environmental variation, while co-localised QTL for different traits may reflect shared developmental or regulatory pathways. Chromosome 7B had the highest number of QTL identified across multiple stomatal anatomical traits and included several closely positioned QTL candidates across sowing times in S1, suggesting a genomic region with potentially environmentally robust control of stomatal anatomy. In addition, six groups of QTL candidates for anatomical traits were repeatedly detected in similar chromosomal regions across both seasons; for the same traits across seasons, stable QTL were identified on 2B for SA, 2B and 5A for GCL, and 7A for GCW (Table S5). Beyond stability across seasons, co-localised QTL were observed where different but closely linked traits mapped to the same region across seasons, including chromosome 5B (SA in S1 and GCL in S2) and 4B (*g_smax_* in S1 and SA in S2). In addition to cross-season co-localisation, a substantial number of QTL exhibiting pleiotropic effects for multiple traits were identified within individual seasons (Tables S4; S5 & S8). Together, these findings suggest that certain genomic regions exert consistent coordinated control over stomatal anatomy and function, either through pleiotropic genes, or tightly linked loci influencing multiple stomatal traits. Identical markers detected across environments on chromosomes 1B and 3A for SA, and on chromosome 7D for GCL, further support the presence of environmentally stable candidate regions (Table S4). Future directions regarding these candidate QTLs and their translation into breeding programs are discussed later.

### Integrated Stomatal Conductance C Anatomy Traits and QTL

*g_se_*, a measure of how closely *g_sop_* operates to its *g_smax_*, was significantly influenced by TOS in S1, genotype in S2 and by leaf surface in both years, highlighting that *g_se_* is shaped by the intrinsic physiological capacity of genotypes and their plasticity in stomatal behaviour under environmental variation. Higher *g_se_* values at TOS 1 across both leaf surfaces in S1 suggest that earlier-sown plants operate closer to their anatomical gas exchange potential under cooler, less stressful conditions. In contrast, the 27.5% decline in *g_se_* at later sowing times in S1 points to heat stress impairing stomatal function, resulting in suboptimal gas exchange despite maintained anatomical capacity. However, no such difference between TOS was detected in S2, suggesting that the effect of sowing time, and thus heat stress, on *g_se_* may not be uniform across seasons. Such divergence could reflect differences in environmental conditions as previously discussed, genotype performance, or their interaction and highlights the complexity of predicting physiological responses to heat. The patterns we observed, particularly in S1, highlight a decoupling of physiological function (*g_sop_*) from structural potential (*g_smax_*) under stress, aligning with prior work by McAusland et al., (2016) and Ochoa et al., (2024), showing that stomata may exhibit delayed or incomplete opening, even when anatomical capacity remains unchanged, leading to inefficient photosynthetic regulation.

Notably, significant genotype-level variation in *g_se_* and its interaction with TOS reveals that some varieties possess superior stomatal regulatory mechanisms, allowing them to maintain efficient gas exchange even under stress. This plasticity in *g_se_* represents a key physiological component of heat resilience and is emerging as a critical trait for breeding heat-resilient wheat. Consistent with this interpretation, *g_se_* exhibited moderate heritability, with higher estimates on the abaxial surface, and all identified QTL candidates for *g_se_* were also located abaxially. Although these QTLs were detected in a single season, their identification suggests that variation in stomatal regulatory efficiency is, at least in part, under genetic control.

Unlike QTLs for anatomical traits, which tend to be more stable across environments, *g_se_*-associated loci appear to be more context-dependent, reflecting the sensitivity of stomatal regulation to environmental conditions. Consequently, further validation across seasons and environments is required before these loci can be confidently deployed in breeding programs.

Together, these findings position *g_se_* as an emerging, higher-order functional trait that bridges anatomical potential and physiological performance, emphasising the need to consider both stomatal anatomy and physiological responsiveness when assessing stomatal traits. Increases in *g_smax_* alongside declines in *g_se_* highlight that selection for genotypes with high *g_smax_* alone is inadequate, and breeding efforts should also account for how efficiently this anatomical capacity is utilised under stress conditions. The identification of genotypes with high *g_se_*, those operating closer to *g_smax_* under stress, along with heritable variation and putative QTL for *g_se_*, highlights a promising, novel breeding target linked to physiological stress resilience. Thus, while selecting for optimal anatomical traits remains valuable, breeding efforts must target genotypes that sustain high operating efficiency under abiotic stress through enhanced stomatal control (Franks et al., 2009).

### Implications for Grain Yield

Grain yield was significantly higher under timely sown conditions compared to delayed sowing in both seasons, and genotypic variation in yield was evident. Importantly, the significant TOS × Genotype interaction indicates that sowing time affects yield differently across genotypes. This suggests that the optimal genotype for yield is not universal but rather context dependent. The ideal combination of stomatal anatomical traits to maximise yield remains unresolved, with some studies linking higher SD and greater *g_smax_* to improved yield and performance, while others report negative relationships between SD or *g_smax_* and yield (McAusland et al., 2021; Samantara et al., 2025). Additionally, other studies have found no correlations between SD and yield, thus warranting further investigation (Liao et al., 2005; Shahinnia et al., 2016). Genotypes that maintain higher *g*ₛ and *g_se_* under later sowing conditions, for example, may have a selective advantage as climate change pushes sowing windows later or increases the frequency of heatwaves during critical periods. These results highlight the importance of selecting genotypes that not only possess favourable stomatal traits but also exhibit resilience to environmental stresses associated with different sowing times.

### Limitations G Future Directions

We recognise some inherent limitations in our study that we address here and highlight areas of future research. This study was conducted at a single location and focused on one developmental stage. Given stomatal anatomical and physiological traits are known to exhibit strong plasticity across developmental stages and in response to local environmental drivers (Balla et al., 2019; Buckley, 2019; Lawson and Blatt, 2014; Li et al., 2025), this constrains the generalisability of the findings across diverse environments and growth phases. Additionally, all measurements were performed under well-watered conditions, and stomatal behaviour under water-limited systems may diverge substantially, particularly the relationship between stomatal traits and yield (Li et al., 2025; Pantha et al., 2025). Under water stress, sustained reductions in stomatal conductance are likely to impose stomatal limitations to photosynthesis that exceed biochemical limitations across the season when compared with heat stress, thereby strengthening links between stomatal traits and yield (Blum, 2009; Richards et al., 2010). In this study, heat stress was imposed via altered sowing time meaning treatments inherently differed in photoperiod. Given stomatal conductance and daily stomatal rhythms in wheat are responsive to day length, some observed differences between sowing times may reflect indirect photoperiod effects in addition to those of temperature stress (Dodd et al., 2005; Yanovsky and Kay, 2001). It is also important to note that the strength of the second season’s results was reduced by the smaller number of genotypes compared to S1, reflected in lower LOD scores for QTL detection. Reduced population size lowers statistical power for detecting loci of small to moderate effect and limits the resolution of QTL confidence intervals, particularly for traits with complex genetic control (Holland, 2007; Xu, 2003). Furthermore, while we have inferred that stomatal size and density correspond to stomatal responsiveness, the relationships between anatomical traits and dynamic stomatal kinetics is context-dependent and influenced by numerous factors (Lawson and Vialet-Chabrand, 2019; McAusland et al., 2016), thus, direct measurements of stomatal kinetics in response to environmental fluctuations remain necessary to validate these assumptions.

To overcome these limitations, future research should expand phenotyping efforts to multiple environments and developmental stages to capture the full spectrum of stomatal behaviour under variable field conditions. Scaling up stomatal trait assessment will benefit from high-throughput approaches, including UAV-based remote sensing such as hyperspectral and thermal imaging and AI-driven analytics, which can provide scalable and precise measurements of stomatal function (Cheng et al., 2024; El-Hendawy et al., 2019; Sadeh et al., 2025). In particular, deeper investigation into stomatal kinetics and their coordination with photosynthetic efficiency and yield performance will refine trait selection for breeding. Novel targeted breeding and genome editing technologies such as Clustered-Regularly-Interspaced-Short-Palindromic-Repeats (CRISPR)-Cas9 efficiently generate precise modifications within a single generation, showing promise in the development of climate resilient wheat (Abdallah et al., 2025; Dunn et al., 2019). Emerging frameworks such as the leveraging multi-omics approaches combined with cutting edge AI-modelling to engineer stomatal regulatory networks also present opportunities to develop stomatal ideotype plants, highlighting innovative directions for integrating anatomical and dynamic stomatal traits into breeding pipelines (Chaplin et al., 2025b). With respect to candidate QTLs, future studies could experimentally verify these QTLs and their potential though fine-mapping.

Alternatively, as stomatal anatomical and physiological traits appear to be linked to numerous QTLs across the genome (Table S4), breeding programs may benefit from developing an extensive training population and using genomic selection for predicting favourable stomatal traits (Meuwissen et al., 2001).

### Conclusions

This study advances our understanding of how stomatal traits are integral to wheat adaptation to different sowing times and, in turn, environmental conditions. By disentangling the effects of *g_s_*, stomatal anatomy and leaf surface, along with quantifying heritability and mapping QTL, we demonstrate that early sowing supports more optimal stomatal function and that genotypes vary significantly in their ability to maintain efficient conductance under thermal stress.

Our results provide clear field-based evidence of environmentally responsive, context-dependent, and genetically controlled stomatal behaviour. 125 putative QTLs we identified across seven stomatal traits, 42 of which were consistent with previously reported QTL for stomatal traits in wheat. Moderate heritability and season-specific, abaxial QTL for *g_s_* further supports a genetic basis for *g_s_* variation. Our results support hypotheses proposing the adaptive value of small, dense stomata for rapid and responsive gas exchange control under stress, yet the ideal combination of anatomical traits for maximising yield remains unresolved and is likely highly context-specific, and may be linked to other environmental factors, including water availability or evaporative demand. Stomatal anatomical traits had higher heritability than conductance traits with 115 candidate QTLs and chromosomal regions for anatomical traits identified, particularly on the abaxial surface. Chromosomes 2B and 5B were identified as promising stable genomic regions controlling stomatal traits, aligned with prior work. 7B also emerges as a strong candidate chromosomal region associated with a high number of stomatal anatomical traits. Six groups of QTLs were also identified in similar chromosomal regions for anatomical traits across both seasons including stable and co-localised pleiotropic loci. These findings emphasise the potential for targeted selection of genetically stable stomatal anatomical traits. The contrasting heritability and environmental stability of anatomical versus conductance traits further underscore the need for environment-specific selection strategies.

Our findings reinforce the importance of integrating structural traits (SA and SD) with functional traits (*g_s_, g_se_*) as a powerful strategy for developing climate-resilient wheat varieties. Collectively, this work provides field-based evidence that stomatal traits exhibit exploitable genetic variation and genomic associations, supporting their inclusion in breeding pipelines aimed at improving wheat performance under increasing heat stress.

## Author Contributions

WS & EC: Conceptualisation; EC, WS & BS: Data Curation; EC & ET: Formal Analysis; WS: Funding Acquisition; EC & WS: Investigation; EC & WS: Methodology; WS: Project Administration; WS: Resources; WS: Software; AM & WS: Supervision; EC, WS & ET: Validation; EC & ET: Visualisation; EC: Writing - Original Draft; EC, AM, WS, ET, BS & RT: Writing – Review & Editing.

## Acknowledgements

We thank colleagues Fiona Foster, Annette Tredrea and Dr Rebecca Thistlethwaite for supporting the field work in this study. We thank Dr Hannah Robinson and Dr Andrew Bowerman for the imputed genotype dataset.

We acknowledge the use of the facilities, and scientific and technical assistance of the Sydney Informatics Hub and the University of Sydney Node of the Australian Plant Phenomics Network (APPN), which is supported by the Australian Government’s National Collaborative Research Infrastructure Strategy (NCRIS) and the Grains Research and Development Corporation.

We acknowledge the use of artificial intelligence (Microsoft Copilot GPT-5) in the preparation of this manuscript, solely in the final stages to enhance clarity, flow and language. We have rigorously vetted all content edited by AI tools. No content, ideas or substantive bodies of text were generated by AI and the manuscript reflects the original ideas and work of the authors.

## Data Availability

Original datasets are openly available in a publicly accessible repository. The original contributions presented in the study including physiological field data and data from genomic analyses are publicly available and the data can be found using the DOI: 10.5281/zenodo.17925401. Genetic sequencing data is not provided due to commercial sensitivity.

All files for the 3D printed leaf clip, the Python script for stomatal annotation and the files and instructions for installing and using the FieldDino App are provided in a public GitHub repository which guides users through each step – https://github.com/williamtsalter/FieldDinoMicroscopy.

## Supplementary Materials

**Figure S1: Schematic representation of procedure for using LI-600 porometer to measure abaxial and adaxial surface stomatal conductance.**

**Figure S2: Representative images of stomatal anatomy collected *in situ* using 200x magnification handheld digital microscope in season 1.** (A) adaxial and (B) abaxial raw images collected using microscope. (C) and (D) show image with automatically labelled stomata with ellipses using the deep learning model we trained.

**Figure S3: Representative images of stomatal anatomy collected *in situ* using 400x magnification handheld digital microscope in season 2.** (A) adaxial and (B) abaxial raw images collected using microscope. (C) and (D) show image with automatically labelled stomata with ellipses using the deep learning model we trained.

**Figure S4: Relationship between stomatal density and stomatal area in season 1.** Shaded markers represent irrigated plants and unshaded markers represent rainfed plants. Circular markers represent abaxial leaf surface and triangular represent adaxial leaf surface.

**Figure S5: LOD score of putative QTLs by trait and chromosome**. The scatterplot shows the LOD scores of putatiative QTLs identified by chromosome and position (in mega base pairs), with each color corresponding to a specific year, TOS and surface.

**Table S1: Overview of germplasm screened in season 1 GRDC field trials.**

**Table S2: Overview of germplasm screened in season 2 of GRDC field trials, selected from the broader University of Sydney genetics program. All lines grown in season 2 were grown in season 1.**

**Table S3: Season 1 (2023) data showing average values for key traits at TOS 1 and at TOS 2, and % change from TOS 1 to TOS 2. Values are given for each surface independently and for both surfaces averaged.**

**Table S4: The number of distinct putative QTL candidates across all trials by trait and chromosome. The “.” represents 0 and if the number is marked with a * then one of the QTL candidates was also detected in another trial.**

**Table S5: List of putative QTL candidates for each trait, organized by trial (indexed by Year and TOS) and surface. The chromosome, position (in base pairs), effect size and LOD score for each corresponding QTL are also provided.**

**Table S6: Season 2 (2024) data showing average values for key traits at TOS 1 and at TOS 2, and % change from TOS 1 to TOS 2. Values are given for each surface independently and for both surfaces averaged.**

**Table S7: The number of QTLs reported in literature by trait and chromosome. The last column shows the number of putative QTLs we found for the same trait and chromosome.**

**Table S8: The number of putative QTLs within a 10 Mbp region by chromosome and trait. The 20 bolded rows indicate a region that may contain pleiotropic QTLs for stomatal traits.**

## Abbreviations

CAIGE: CIMMYT Australia ICARDA Germplasm Evaluation
CIMMYT: International Maize and Wheat Improvement Center
CO_2_: Carbon Dioxide
CRISPR: Clustered-Regularly-Interspaced-Short-Palindromic-Repeats
EDPIE: Elite Diversity International Experiment
ESWYT: Elite Selection Wheat Yield Trial
GCL: Guard Cell Length
GCW: Guard Cell Width
*g_s_*: Stomatal conductance
*g_se_*: *g_sop_/g_smax_* - Stomatal conductance operating efficiency (unitless)
*g_smax_*: Maximum anatomical stomatal conductance
*g_sop_*: Operating rate of stomatal conductance
H_2_O: Water
HTWYT: High Temperature Wheat Yield Trials
ICARDA: International Centre for Agricultural Research in the Dry Areas
IRGA: Infra-red gas analysers
mAP: Mean average precision
Mbp: Mega base pair
LOD: Logarithm of the odds
PSII: Photosystem II
QTL: Quantitative Trait Loci
SATYN: Stress Adapted Trait Yield Nursery
SAWYT: Semi-Arid Wheat Yield Trial
S1: Season 1; 2023
S2: Season 2; 2024
SA: Stomatal Area
SD: Stomatal density (stomata m^-2^)
TKW: Thousand Kernel Weight
TOS: Time of Sowing
VPD: Vapour Pressure Deficit
WUE: Water Use Efficiency
YOLO: You Only Look Once

