## Supplementary Materials for "QTL for Heat-Induced Stomatal Anatomy Underpin Gas Exchange Variation in Field-Grown Wheat"

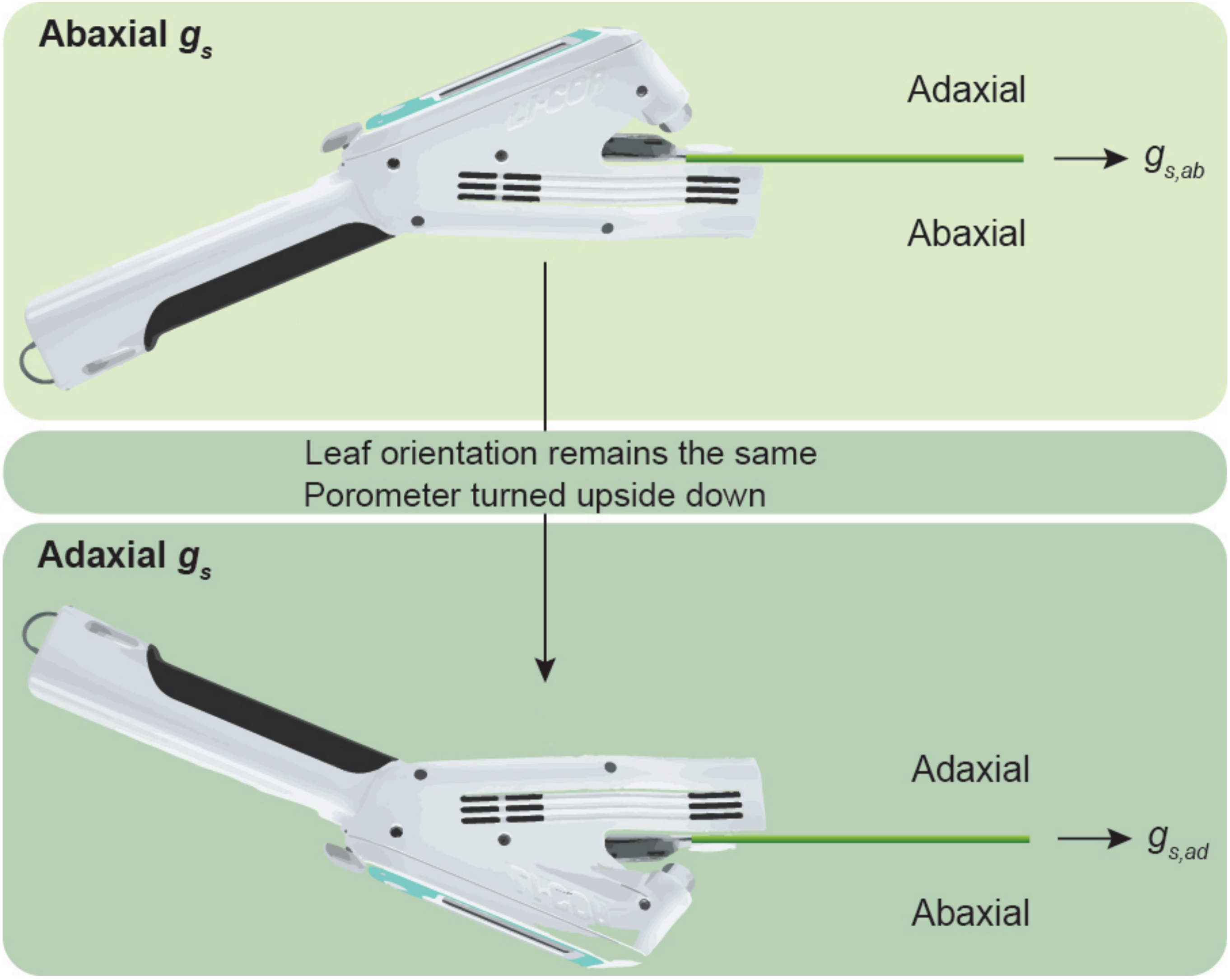


**Figure S1: Schematic representation of procedure for using LI-600 porometer to measure abaxial and adaxial surface stomatal conductance.**

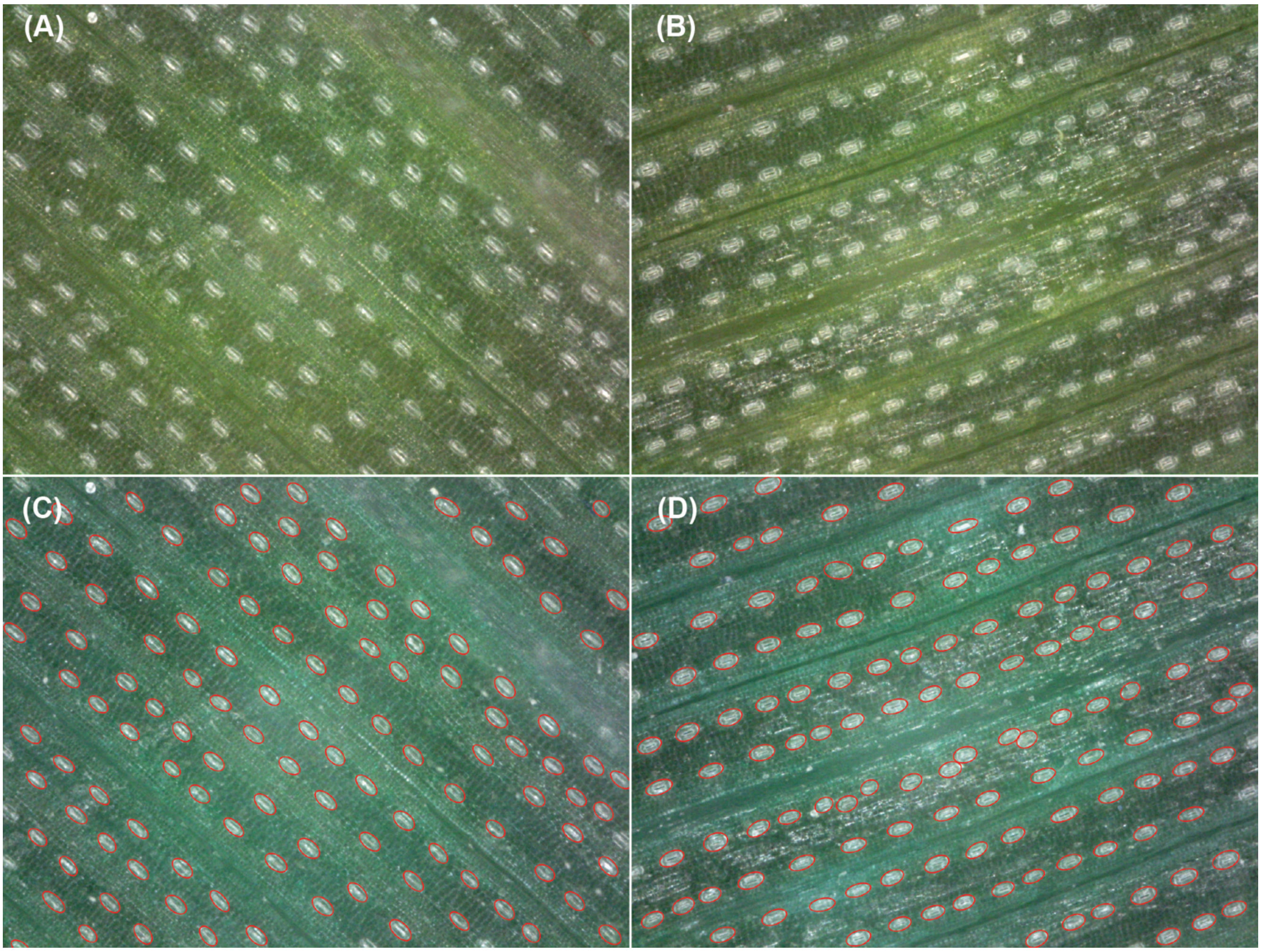


**Figure S2: Representative images of stomatal anatomy collected *in situ* using 200x magnification handheld digital microscope in season 1.** (A) adaxial and (B) abaxial raw images collected using microscope. (C) and (D) show image with automatically labelled stomata with ellipses using the deep learning model we trained.

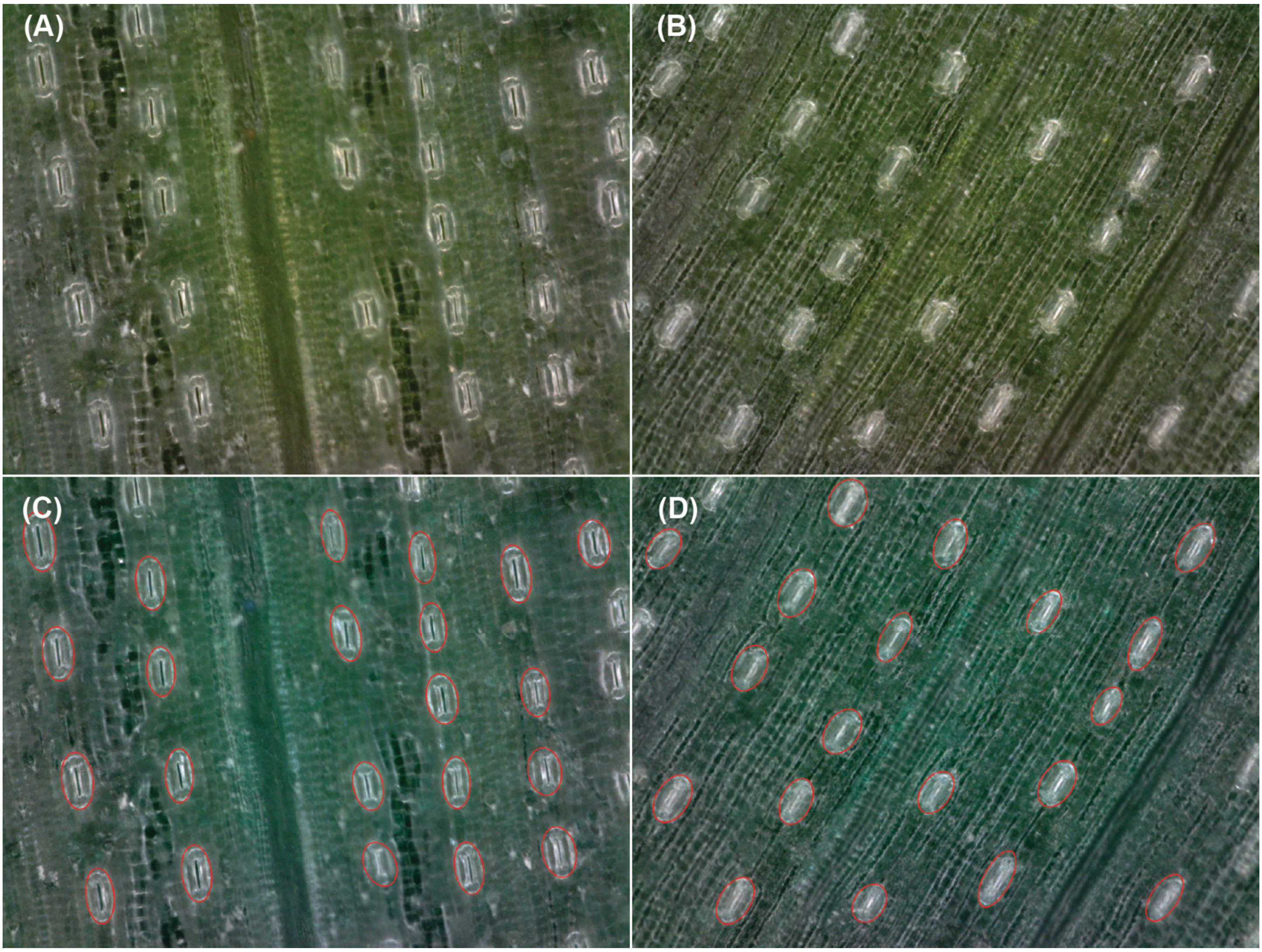


**Figure S3: Representative images of stomatal anatomy collected *in situ* using 400x magnification handheld digital microscope in season 2.** (A) adaxial and (B) abaxial raw images collected using microscope. (C) and (D) show image with automatically labelled stomata with ellipses using the deep learning model we trained.

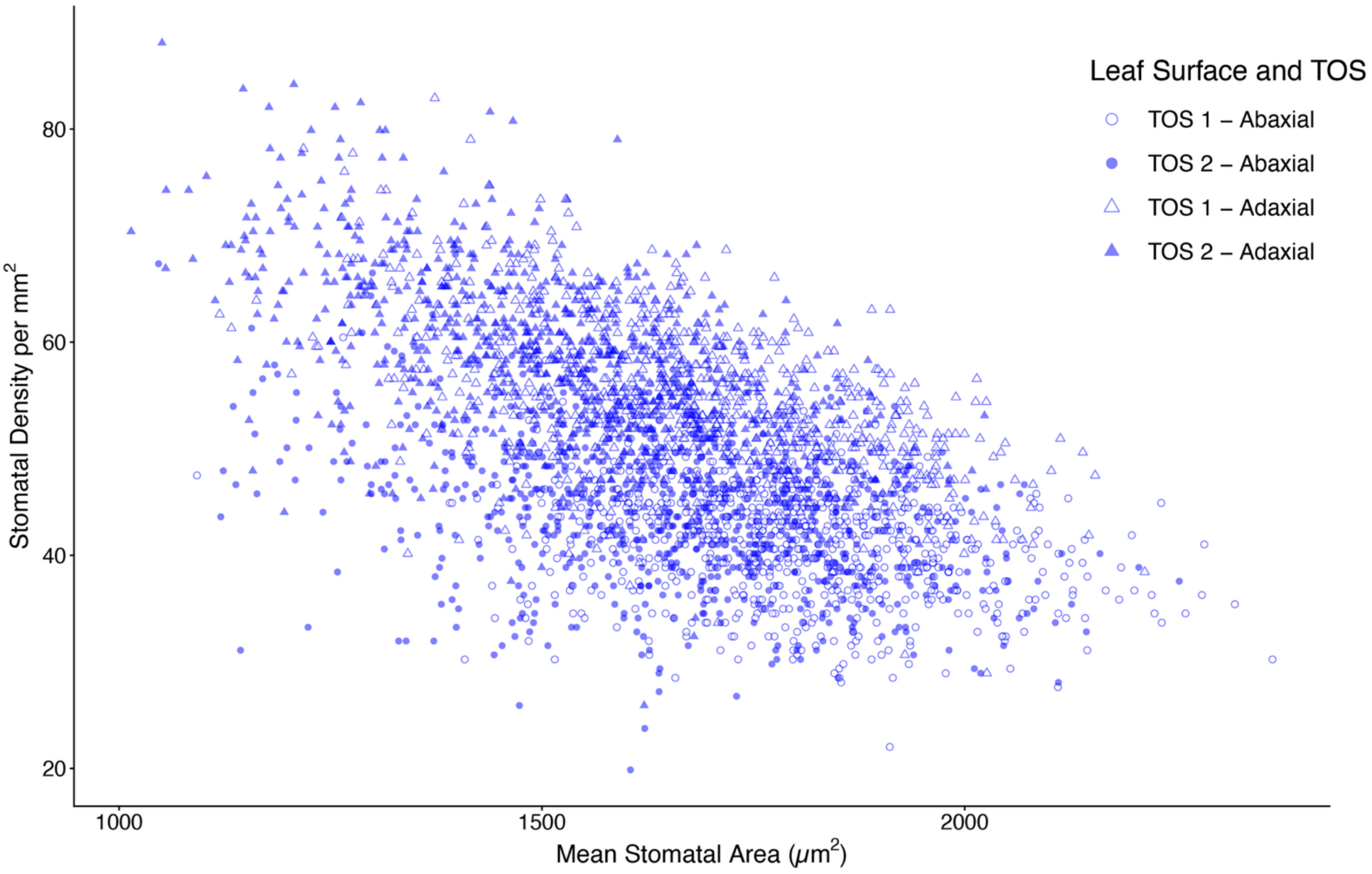


**Figure S4: Relationship between stomatal density and stomatal area in season 1.** Shaded markers represent irrigated plants and unshaded markers represent rainfed plants. Circular markers represent abaxial leaf surface and triangular represent adaxial leaf surface.

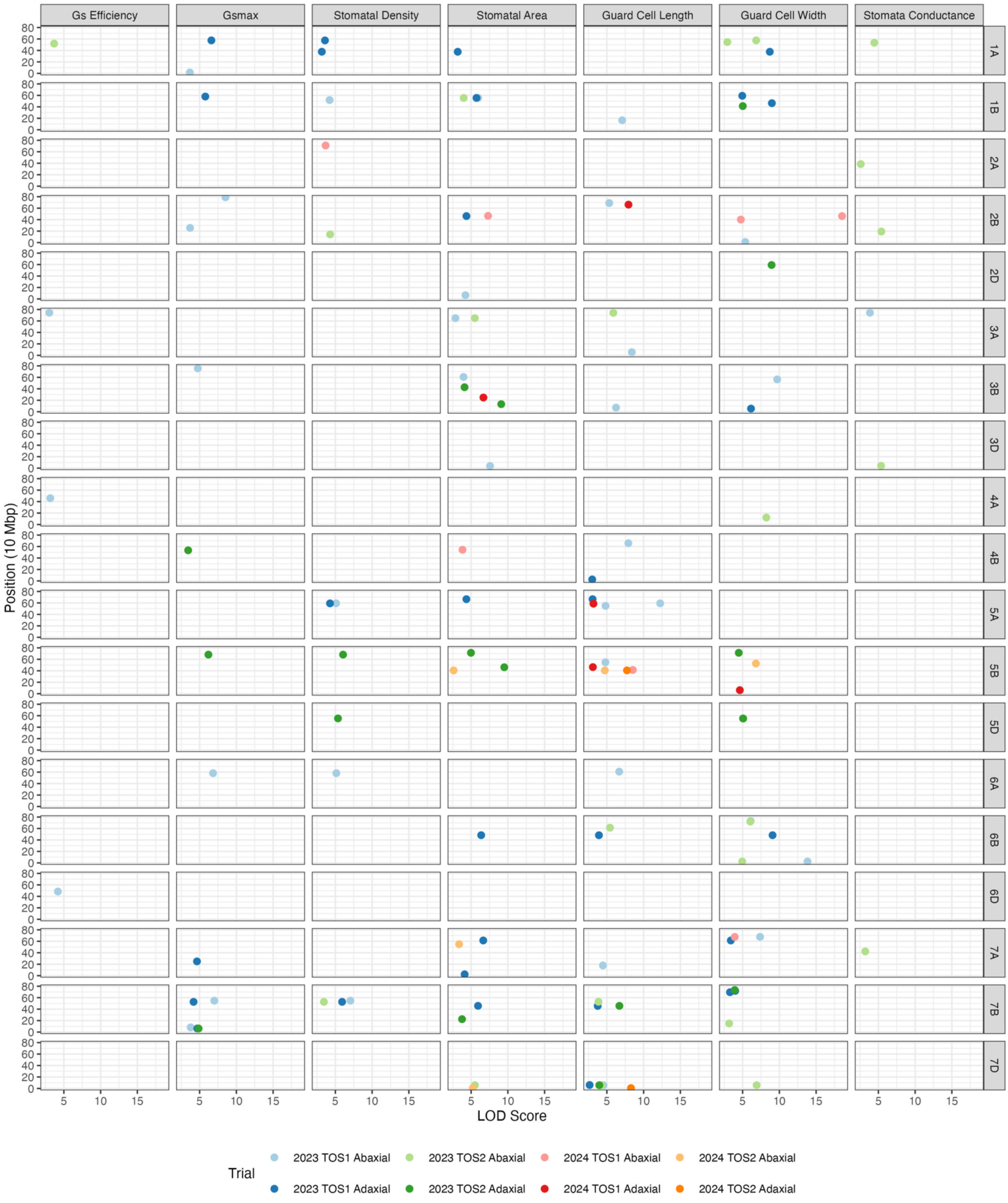


**Figure S5: LOD score of putative QTLs by trait and chromosome**. The scatterplot shows the LOD scores of putatiative QTLs identified by chromosome and position (in mega base pairs), with each color corresponding to a specific year, TOS and surface.

**Table S1: Overview of germplasm screened in season 1 GRDC field trials.**

| No. | Designation | No. | Designation |
|---|---|---|---|
| 1 | PBI09C009-BC-DH80 | 101 | PBI17N014-0C-0N-0N-010N-4N |
| 2 | PBI09C008-BC-DH17 | 102 | PBI17N014-0C-0N-0N-010N-5N |
| 3 | PBI09C043-BC-0C-20N-99N | 103 | PBI17N014-0C-0N-0N-010N-6N |
| 4 | PBI09C048-BC-0C-6N-99N | 104 | PBI17N014-0C-0N-0N-010N-8N |
| 5 | ACIAR09PBI C04-52C-DH3 | 105 | PBI17N014-0C-0N-0N-010N-16N |
| 6 | ACIAR09PBI C04-52C-DH10 | 106 | PBI17N014-0C-0N-0N-010N-19N |
| 7 | PBI07C101-DH96 | 107 | PBI17N015-0C-0N-0N-010N-2N |
| 8 | PBI07C101-DH147 | 108 | PBI17N015-0C-0N-0N-010N-5N |
| 9 | CMSA05Y00954T-040M-040ZTP0Y-040ZTM-040SY-12ZTM-01Y-0B | 109 | PBI17N015-0C-0N-0N-010N-8N |
| 10 | SUNTOP | 110 | PBI17N015-0C-0N-0N-010N-10N |
| 11 | ICW02.00099-11APTS-0AP-0AP-9AP-0AP | 111 | PBI17N015-0C-0N-0N-010N-19N |
| 12 | ICW04.20024-9AP-0AP-0AP-0AP-9AP-0AP | 112 | PBI17N015-0C-0N-0N-010N-24N |
| 13 | CMSS08B00645T-099TOPY-099M-099NJ-099NJ-6WGY-0B | 113 | PBI17N016-0C-0N-0N-010N-27N |
| 14 | CMSS08B00648T-099TOPY-099M-099NJ-2WGY-0B | 114 | PBI17N018-0C-0N-0N-010N-1N |
| 15 | CMSA08Y00613S-050Y-050ZTM-050Y-59BMX-010Y-0B | 115 | PBI17N019-0C-0N-0N-010N-6N |
| 16 | CMSA08M00007T-030(AWW5L7BO1HET)Y-040M-0NJ-4Y-0B | 116 | PBI17N020-0C-0N-0N-010N-12N |
| 17 | SUNLIN | 117 | PBI17N021-0C-0N-0N-010N-5N |
| 18 | CMSS06Y00885T-099TOPM-099Y-099ZTM-099NJ-099NJ-26WGY-0B | 118 | PBI16C001-0C-0N-0N-010N-5N |
| 19 | CMSA06M00008T-024(PINBD1BHET)Y-040ZTM-026(PINBD1BPOS)ZTY-20ZTM-0Y-0B | 119 | PBI16C007-0C-0N-0N-010N-2N |
| 20 | CMSA05M00140S-021(CRE1M19-GWM577BO1)M-028(BO1 POS& CRE1 POS)ZTY-040ZTM-040SY-11ZTM-0Y-0B | 120 | PBI16C008-0C-0N-0N-010N-7N |
| 21 | MACE | 121 | PBI16C009-0C-0N-0N-010N-14N |
| 22 | SCOUT | 122 | PBI16C009-0C-0N-0N-010N-15N |
| 23 | CMSA05Y01186T-040M-040ZTP0Y-040ZTM-040SY-32ZTM-02Y-0B | 123 | PBI16C009-0C-0N-0N-010N-17N |
| 24 | CGSS05B00258T-099TOPY-099M-099Y-099ZTM-12WGY-0B | 124 | PBI16C010-0C-0N-0N-010N-6N |
| 25 | SUNCO | 125 | PBI16C011-0C-0N-0N-010N-1N |
| 26 | BIOINTA 1005 | 126 | PBI16C011-0C-0N-0N-010N-5N |

| | | | |
|---|---|---|---|
| 27 | BORLAUG 100 | 127 | PBI16C011-0C-0N-0N-010N-10N |
| 28 | COOLAH | 128 | PBI16C013-0C-0N-0N-010N-4N |
| 29 | CUTLASS | 129 | PBI16C013-0C-0N-0N-010N-12N |
| 30 | SCEPTER | 130 | PBI16C015-0C-0N-0N-010N-10N |
| 31 | CONDO | 131 | PBI16C015-0C-0N-0N-010N-13N |
| 32 | FLANKER | 132 | PBI16C016-0C-0N-0N-010N-11N |
| 33 | VIKING | 133 | PBI16C016-0C-0N-0N-010N-14N |
| 34 | ICW05.0660-18AP-0AP-0AP-1AP-0AP | 134 | TUK004-0Q-0N-020N-0N-5N |
| 35 | CMSS11B00167S-099M-0SY-11M-0WGY | 135 | TUK004-0Q-0N-020N-0N-8N |
| 36 | PTSS02Y00021S-099B-099Y-030ZTM-040SY-040M-21Y-0M-0SY-0Y-0Y | 136 | TUK004-0Q-0N-020N-0N-36N |
| 37 | PTSS02Y00023S-099B-099Y-099B-099Y-100B-0Y | 137 | PTSS15Y00010S-099B-099Y-099M-2Y-020Y |
| 38 | PTSS08GHB00013S-0SH-099SHB-099SHB-099SHB-32Y-0Y | 138 | PTSS15Y00023S-099B-099Y-099M-23Y-020Y |
| 39 | PTSS08GHB00016S-099SH-099Y-099SHB-099Y-155Y-0Y | 139 | PTSS15Y00024S-099B-099Y-099M-2Y-020Y |
| 40 | SDSS12B00849T-0Y-0B-0B-12Y-0B | 140 | PTSS15Y00024S-099B-099Y-099M-10Y-020Y |
| 41 | SUNCHASER | 141 | PTSS15Y00024S-099B-099Y-099M-17Y-020Y |
| 42 | VIXEN | 142 | PTSS14Y00062S-0B-099Y-099B-39Y-020Y |
| 43 | IMPALA | 143 | PTSS14Y00071S-0B-099Y-099B-33Y-020Y |
| 44 | RELIANT | 144 | PTSS14Y00328S-0B-099Y-099B-32Y-020Y |
| 45 | BECKOM | 145 | PTSS14Y00329S-0B-099Y-099B-33Y-020Y |
| 46 | HELLFIRE | 146 | PTSS11Y00152S-0SHB-099B-099Y-099B-099Y-17Y-020Y-0B |
| 47 | MUSTANG | 147 | PTSS15Y00053S-099B-099Y-099M-8Y-020Y-0B |
| 48 | PBIC15039-0C-9N-010N-2N-0N | 148 | PTSS15Y00056S-099B-099Y-099M-14Y-020Y-0B |
| 49 | PBIC15038-0C-29N-010N-1N-0N | 149 | PTSS15Y00082S-099B-099Y-099M-16Y-020Y-0B |
| 50 | PBIC15034-0C-24N-010N-1N-0N | 150 | PTSS02B00102T-0TOPY-0B-0Y-0B-11Y-0M-0SY-0B-0Y-4Y-0M |
| 51 | PBIC15034-0C-63N-010N-1N-0N | 151 | PTSS14Y00330S-0B-099Y-099B-40Y-0B |
| 52 | PBIC15034-0C-73N-010N-1N-0N | 152 | PTSS14Y00057S-0B-099Y-099B-10Y-020Y-0B |

| | | | |
|---|---|---|---|
| 53 | PBIC15030-0C-13N-010N-1N-0N | 153 | PTSS14Y00013S-0B-099Y-099B-14Y-020Y-0B |
| 54 | PBI19N002-0N-5N | 154 | PTSS15Y00032S-099B-099Y-099M-25Y-020Y |
| 55 | PBI19N003-0N-11N | 155 | PTSS15Y00050S-099B-099Y-099M-1Y-020Y |
| 56 | PBI19N007-0N-3N | 156 | CATAPULT |
| 57 | PBI19N007-0N-6N | 157 | SUNMASTER |
| 58 | PBI19N007-0N-8N | 159 | ROCKSTAR |
| 59 | PBI17N001-0N-0N-12N | 158 | SHERIFF CL PLUS |
| 60 | PBI17N002-0N-0N-38N | 160 | VALIANT CL PLUS |
| 61 | PBI17N002-0N-0N-76N | 161 | HAVOC |
| 62 | PBI17N003-0N-0N-32N | 162 | STEALTH |
| 63 | PBI17N004-0N-0N-16N | 163 | IG-ANU-HeatLine-001 |
| 64 | PBI17N004-0N-0N-18N | 164 | IG-ANU-HeatLine-002 |
| 65 | PBI17N005-0N-0N-29N | 165 | IG-ANU-HeatLine-008 |
| 66 | PBI17N005-0N-0N-32N | 166 | IG-ANU-HeatLine-009 |
| 67 | PBI17N006-0N-0N-89N | 167 | IG-ANU-HeatLine-011 |
| 68 | PBI17N006-0N-0N-96N | 168 | IG-ANU-HeatLine-023 |
| 69 | PBI17N007-0N-0N-13N | 169 | IG-ANU-HeatLine-028 |
| 70 | PBI17N008-0N-0N-10N | 170 | IG-ANU-HeatLine-033 |
| 71 | PBI17N008-0N-0N-113N | 171 | IG-ANU-HeatLine-034 |
| 72 | PBI17N009-0N-0N-4N | 172 | IG-ANU-HeatLine-038 |
| 73 | PBI17N009-0N-0N-46N | 173 | IG-ANU-HeatLine-055 |
| 74 | PBI17N011-0N-0N-20N | 174 | IG-ANU-HeatLine-056 |
| 75 | PBI17N011-0N-0N-41N | 175 | IG-ANU-HeatLine-074 |
| 76 | PBI17N012-0N-0N-106N | 176 | IG-ANU-HeatLine-078 |
| 77 | PBI17N013-0N-0N-3N | 177 | IG-ANU-HeatLine-080 |
| 78 | PBI17N013-0N-0N-62N | 178 | IG-ANU-HeatLine-081 |
| 79 | PBI17N014-0N-0N-54N | 179 | IG-ANU-HeatLine-088 |
| 80 | PBI17N015-0N-0N-58N | 180 | IG-ANU-HeatLine-097 |
| 81 | PBI17N015-0N-0N-96N | 181 | IG-ANU-HeatLine-103 |
| 82 | PBI17N001-0C-0N-0N-010N-8N | 182 | IG-ANU-HeatLine-107 |
| 83 | PBI17N002-0C-0N-0N-010N-16N | 183 | IG-ANU-HeatLine-010 |
| 84 | PBI17N004-0C-0N-0N-010N-5N | 184 | IG-ANU-HeatLine-025 |
| 85 | PBI17N005-0C-0N-0N-010N-8N | 185 | IG-ANU-HeatLine-026 |
| 86 | PBI17N005-0C-0N-0N-010N-10N | 186 | IG-ANU-HeatLine-035 |
| 87 | PBI17N005-0C-0N-0N-010N-14N | 187 | IG-ANU-HeatLine-037 |
| 88 | PBI17N006-0C-0N-0N-010N-3N | 188 | IG-ANU-HeatLine-047 |

| | | | |
|---|---|---|---|
| 89 | PBI17N007-0C-0N-0N-010N-8N | 189 | IG-ANU-HeatLine-051 |
| 90 | PBI17N007-0C-0N-0N-010N-11N | 190 | IG-ANU-HeatLine-059 |
| 91 | PBI17N007-0C-0N-0N-010N-15N | 191 | IG-ANU-HeatLine-063 |
| 92 | PBI17N009-0C-0N-0N-010N-10N | 192 | IG-ANU-HeatLine-067 |
| 93 | PBI17N010-0C-0N-0N-010N-5N | 193 | SHAMIEKH-3 |
| 94 | PBI17N010-0C-0N-0N-010N-6N | 194 | ALATHEER-4 |
| 95 | PBI17N010-0C-0N-0N-010N-10N | 195 | ISR 812.8/CARINYA |
| 96 | PBI17N010-0C-0N-0N-010N-13N | 196 | CMSS08B00648T-099TOPY-099M-099NJ-2WGY-0B |
| 97 | PBI17N012-0C-0N-0N-010N-17N | 197 | PBI16C001-0C-0N-0N-010N-12N |
| 98 | PBI17N013-0C-0N-0N-010N-7N | 198 | CMSA05Y00954T-040M-040ZTP0Y-040ZTM-040SY-12ZTM-01Y-0B |
| 99 | PBI17N013-0C-0N-0N-010N-8N | 199 | PTSS02B00096T-0TOPY-0B-0Y-0B-33Y-0M-0SY-0Y-0Y |
| 100 | PBI17N013-0C-0N-0N-010N-9N | 200 | PTSS02B00094T-0TOPY-0B-0Y-0B-3Y-0ZTB-0SY-0Y-0Y |

**Table S2: Overview of germplasm screened in season 2 of GRDC field trials, selected from the broader University of Sydney genetics program. All lines grown in season 2 were grown in season 1.**

| No. | Designation | No. | Designation |
|---|---|---|---|
| 1 | Borlaug 100 | 26 | PBI17N002-0C-0N-0N-010N-16N |
| 2 | Cutlass | 27 | PBI17N005-0N-0N-32N |
| 3 | Hellfire | 28 | PBI17N006-0N-0N-89N |
| 4 | IG-ANU-HeatLine-008 | 29 | PBI17N007-0C-0N-0N-010N-8N |
| 5 | IG-ANU-HeatLine-009 | 30 | PBI17N007-0N-0N-13N |
| 6 | IG-ANU-HeatLine-010 | 31 | PBI17N009-0N-0N-46N |
| 7 | IG-ANU-HeatLine-011 | 32 | PBI17N011-0N-0N-20N |
| 8 | IG-ANU-HeatLine-023 | 33 | PBI17N013-0C-0N-0N-010N-8N |
| 9 | IG-ANU-HeatLine-037 | 34 | PBI17N014-0C-0N-0N-010N-19N |
| 10 | IG-ANU-HeatLine-038 | 35 | PBI17N015-0C-0N-0N-010N-10N |
| 11 | IG-ANU-HeatLine-047 | 36 | PBI17N015-0C-0N-0N-010N-8N |
| 12 | IG-ANU-HeatLine-051 | 37 | PBI17N015-0N-0N-96N |
| 13 | IG-ANU-HeatLine-055 | 38 | PBI17N016-0C-0N-0N-010N-27N |
| 14 | IG-ANU-HeatLine-056 | 39 | PBI17N019-0C-0N-0N-010N-6N |
| 15 | IG-ANU-HeatLine-059 | 40 | PBI19N007-0N-3N |
| 16 | IG-ANU-HeatLine-063 | 41 | PBI19N007-0N-8N |
| 17 | IG-ANU-HeatLine-103 | 42 | PBIC15034-0C-24N-010N-1N-0N |
| 18 | IG-ANU-HeatLine-107 | 43 | Reliant |
| 19 | Mace | 44 | RockStar |
| 20 | Mustang | 45 | Scepter |
| 21 | PBI07C101-DH96 | 46 | Scout |
| 22 | PBI16C009-0C-0N-0N-010N-14N | 47 | Stealth |
| 23 | PBI16C013-0C-0N-0N-010N-4N | 48 | Sunchaser |
| 24 | PBI16C016-0C-0N-0N-010N-14N | 49 | Sunco |
| 25 | PBI17N001-0C-0N-0N-010N-8N | 50 | Suntop |

**Table S3: Season 1 (2023) data showing average values for key traits at TOS 1 and at TOS 2, and % change from TOS 1 to TOS 2. Values are given for each surface independently and for both surfaces averaged.**

| Trait & Surface | | TOS 1 | TOS 2 | % Change TOS 1 to TOS 2 |
|---|---|---|---|---|
| **Abaxial** | | | | |
| $g_s$ | mol.m$^{-2}$.s$^{-1}$ | 0.065 | 0.055 | -14.77 |
| **Stomatal Density** | per mm$^{-2}$ | 41.58 | 44.23 | 6.37 |
| **Guard cell Length** | µm | 64.73 | 62.19 | -3.93 |
| **Guard Cell Width** | µm | 39.62 | 38.44 | -2.99 |
| **Stomatal Area** | µm$^{-2}$ | 1785.90 | 1658.94 | -7.11 |
| $g_{smax}$ | mol.m$^{-2}$.s$^{-1}$ | 1.13 | 1.16 | 2.11 |
| $g_{se}$ | - | 0.06 | 0.05 | -15.25 |
| **Adaxial** | | | | |
| $g_s$ | mol.m$^{-2}$.s$^{-1}$ | 0.186 | 0.123 | -33.89 |
| **Stomatal Density** | per mm$^{-2}$ | 54.84 | 58.52 | 6.70 |
| **Guard cell Length** | µm | 65.86 | 61.81 | -6.14 |
| **Guard Cell Width** | µm | 36.99 | 35.91 | -2.91 |
| **Stomatal Area** | µm$^{-2}$ | 1672.63 | 1516.58 | -9.33 |
| $g_{smax}$ | mol.m$^{-2}$.s$^{-1}$ | 1.52 | 1.52 | 0.06 |
| $g_{se}$ | - | 0.12 | 0.08 | -33.33 |
| **Adaxial & Abaxial Average** | | | | |
| $g_s$ | mol.m$^{-2}$.s$^{-1}$ | 0.125 | 0.089 | -28.93 |
| **Stomatal Density** | per mm$^{-2}$ | 48.21 | 51.37 | 6.56 |
| **Guard cell Length** | µm | 65.30 | 62.00 | -5.05 |
| **Guard Cell Width** | µm | 38.31 | 37.17 | -2.95 |
| **Stomatal Area** | µm$^{-2}$ | 1729.27 | 1587.76 | -8.18 |
| $g_{smax}$ | mol.m$^{-2}$.s$^{-1}$ | 1.33 | 1.34 | 0.93 |
| $g_{se}$ | - | 0.09 | 0.07 | -27.53 |
| **Yield** | t.ha$^{-1}$ | 4.62 | 2.79 | -39.64 |

**Table S4: The number of distinct putative QTL candidates across all trials by trait and chromosome. The "." represents 0 and if the number is marked with a * then one of the QTL candidates was also detected in another trial.**

| Chromosome | $g_s$ | GCW | GCL | SA | SD | $g_{smax}$ | $g_{se}$ |
|---|---|---|---|---|---|---|---|
| **1A** | 1 | 3 | . | 1 | 2 | 2 | 1 |
| **1B** | . | 3 | 1 | 3* | 1 | 1 | . |
| **2A** | 1 | . | . | . | 1 | . | . |
| **2B** | 1 | 3 | 2 | 2 | 1 | 2 | . |
| **2D** | . | 1 | . | 1 | . | . | . |
| **3A** | 1 | . | 2 | 2* | . | . | 1 |
| **3B** | . | 2 | 1 | 4 | . | 1 | . |
| **3D** | 1 | . | . | 1 | . | . | . |
| **4A** | . | 1 | . | . | . | . | 1 |
| **4B** | . | . | 2 | 1 | . | 1 | . |
| **5A** | . | . | 4 | 1 | 2 | . | . |
| **5B** | . | 3 | 5 | 3 | 1 | 1 | . |
| **5D** | . | 2 | . | . | 1 | . | . |
| **6A** | . | . | 1 | . | 1 | 1 | . |
| **6B** | . | 5 | 2 | 1 | . | . | . |
| **6D** | . | . | . | . | . | . | 1 |
| **7A** | 1 | 3 | 1 | 3 | . | 1 | . |
| **7B** | . | 4 | 3 | 2 | 3 | 5 | . |
| **7D** | . | 1 | 5* | 2 | . | . | . |
| **Total** | **6** | **31** | **29** | **27** | **13** | **15** | **4** |

**Table S5: List of putative QTL candidates for each trait, organized by trial (indexed by Year and TOS) and surface. The chromosome, position (in base pairs), effect size and LOD score for each corresponding QTL are also provided.**

| Trait | Surface | TOS | Year | Chr | Position (bp) | Size | LOD |
|---|---|---|---|---|---|---|---|
| Guard Cell Length | Abaxial | TOS1 | 2023 | 1B | 165,914,784 | -0.79 | 7.09 |
| Guard Cell Length | Abaxial | TOS1 | 2023 | 2B | 688,831,778 | 1.44 | 5.32 |
| Guard Cell Length | Abaxial | TOS1 | 2023 | 3A | 54,957,133 | -1.92 | 8.40 |
| Guard Cell Length | Abaxial | TOS1 | 2023 | 3B | 71,881,505 | -2.17 | 6.24 |
| Guard Cell Length | Abaxial | TOS1 | 2023 | 4B | 657,893,676 | -0.72 | 7.93 |
| Guard Cell Length | Abaxial | TOS1 | 2023 | 5A | 548,350,881 | -0.76 | 4.81 |
| Guard Cell Length | Abaxial | TOS1 | 2023 | 5A | 591,952,394 | -0.96 | 12.26 |
| Guard Cell Length | Abaxial | TOS1 | 2023 | 5B | 548,494,031 | -0.80 | 4.80 |
| Guard Cell Length | Abaxial | TOS1 | 2023 | 6A | 608,277,937 | -1.76 | 6.68 |
| Guard Cell Length | Abaxial | TOS1 | 2023 | 7A | 178,191,061 | -1.03 | 4.46 |
| Guard Cell Length | Abaxial | TOS1 | 2023 | 7D | 56,579,964 | 0.69 | 4.54 |
| Guard Cell Length | Adaxial | TOS1 | 2023 | 4B | 25,583,788 | -1.24 | 3.01 |
| Guard Cell Length | Adaxial | TOS1 | 2023 | 5A | 665,150,454 | 1.40 | 3.06 |
| Guard Cell Length | Adaxial | TOS1 | 2023 | 6B | 482,341,150 | -0.92 | 3.92 |
| Guard Cell Length | Adaxial | TOS1 | 2023 | 7B | 455,731,677 | -0.83 | 3.75 |

| | | | | | | | |
|---|---|---|---|---|---|---|---|
| Guard Cell Length | Adaxial | TOS1 | 2023 | 7D | 60,474,599 | -0.84 | 2.66 |
| Guard Cell Length | Abaxial | TOS2 | 2023 | 3A | 741,360,511 | -1.58 | 5.88 |
| Guard Cell Length | Abaxial | TOS2 | 2023 | 6B | 613,937,788 | -1.20 | 5.43 |
| Guard Cell Length | Abaxial | TOS2 | 2023 | 7B | 528,725,776 | -1.06 | 3.86 |
| Guard Cell Length | Abaxial | TOS2 | 2023 | 7D | 56,638,565 | 1.09 | 4.27 |
| Guard Cell Length | Adaxial | TOS2 | 2023 | 7B | 457,347,337 | -1.25 | 6.70 |
| Guard Cell Length | Adaxial | TOS2 | 2023 | 7D | 56,638,565 | 1.14 | 3.97 |
| Guard Cell Length | Abaxial | TOS1 | 2024 | 5B | 414,713,237 | 1.86 | 8.54 |
| Guard Cell Length | Adaxial | TOS1 | 2024 | 2B | 661,448,323 | 2.89 | 7.94 |
| Guard Cell Length | Adaxial | TOS1 | 2024 | 5A | 586,828,417 | 1.09 | 3.17 |
| Guard Cell Length | Adaxial | TOS1 | 2024 | 5B | 464,325,673 | -1.22 | 3.10 |
| Guard Cell Length | Abaxial | TOS2 | 2024 | 5B | 406,738,159 | 1.85 | 4.72 |
| Guard Cell Length | Adaxial | TOS2 | 2024 | 5B | 407,109,507 | -1.84 | 7.73 |
| Guard Cell Length | Adaxial | TOS2 | 2024 | 7D | 6,846,653 | 2.24 | 8.28 |
| Guard Cell Width | Abaxial | TOS1 | 2023 | 2B | 9,594,208 | 0.53 | 5.36 |
| Guard Cell Width | Abaxial | TOS1 | 2023 | 3B | 564,634,085 | 0.83 | 9.71 |
| Guard Cell Width | Abaxial | TOS1 | 2023 | 6B | 23,602,849 | -0.96 | 13.83 |

| | | | | | | | |
|---|---|---|---|---|---|---|---|
| Guard Cell Width | Abaxial | TOS1 | 2023 | 6B | 730,602,191 | 0.67 | 6.09 |
| Guard Cell Width | Abaxial | TOS1 | 2023 | 7A | 677,126,838 | -0.76 | 7.37 |
| Guard Cell Width | Adaxial | TOS1 | 2023 | 1A | 376,842,796 | 2.38 | 8.70 |
| Guard Cell Width | Adaxial | TOS1 | 2023 | 1B | 463,960,656 | -1.16 | 8.98 |
| Guard Cell Width | Adaxial | TOS1 | 2023 | 1B | 592,363,151 | -0.63 | 4.95 |
| Guard Cell Width | Adaxial | TOS1 | 2023 | 3B | 54,798,491 | 0.60 | 6.14 |
| Guard Cell Width | Adaxial | TOS1 | 2023 | 6B | 482,975,955 | -0.91 | 9.07 |
| Guard Cell Width | Adaxial | TOS1 | 2023 | 7A | 614,132,821 | 1.13 | 3.39 |
| Guard Cell Width | Adaxial | TOS1 | 2023 | 7B | 695,071,183 | 0.44 | 3.28 |
| Guard Cell Width | Adaxial | TOS1 | 2023 | 7B | 718,926,995 | -0.47 | 3.99 |
| Guard Cell Width | Abaxial | TOS2 | 2023 | 1A | 546,091,806 | -0.42 | 2.92 |
| Guard Cell Width | Abaxial | TOS2 | 2023 | 1A | 575,536,483 | 0.97 | 6.84 |
| Guard Cell Width | Abaxial | TOS2 | 2023 | 4A | 121,156,643 | 1.13 | 8.23 |
| Guard Cell Width | Abaxial | TOS2 | 2023 | 5D | 547,047,053 | 0.65 | 5.11 |
| Guard Cell Width | Abaxial | TOS2 | 2023 | 6B | 23,484,434 | -0.53 | 4.94 |
| Guard Cell Width | Abaxial | TOS2 | 2023 | 6B | 720,739,421 | -0.63 | 6.04 |
| Guard Cell Width | Abaxial | TOS2 | 2023 | 7B | 149,918,869 | 0.97 | 3.17 |

| | | | | | | | |
|---|---|---|---|---|---|---|---|
| Guard Cell Width | Abaxial | TOS2 | 2023 | 7D | 56,579,964 | 0.78 | 6.91 |
| Guard Cell Width | Adaxial | TOS2 | 2023 | 1B | 413,270,635 | -0.48 | 5.02 |
| Guard Cell Width | Adaxial | TOS2 | 2023 | 2D | 591,777,429 | 0.96 | 8.94 |
| Guard Cell Width | Adaxial | TOS2 | 2023 | 5B | 713,297,219 | 0.46 | 4.47 |
| Guard Cell Width | Adaxial | TOS2 | 2023 | 5D | 552,286,738 | 0.51 | 5.06 |
| Guard Cell Width | Adaxial | TOS2 | 2023 | 7B | 733,841,567 | 0.51 | 3.93 |
| Guard Cell Width | Abaxial | TOS1 | 2024 | 2B | 401,211,770 | 1.07 | 4.76 |
| Guard Cell Width | Abaxial | TOS1 | 2024 | 2B | 462,199,334 | -2.16 | 18.53 |
| Guard Cell Width | Abaxial | TOS1 | 2024 | 7A | 675,787,491 | -0.94 | 3.93 |
| Guard Cell Width | Adaxial | TOS1 | 2024 | 5B | 61,842,212 | -1.27 | 4.62 |
| Guard Cell Width | Abaxial | TOS2 | 2024 | 5B | 526,324,523 | 1.54 | 6.81 |
| Stomatal Area | Abaxial | TOS1 | 2023 | 1B | 555,412,597 | 66.39 | 5.95 |
| Stomatal Area | Abaxial | TOS1 | 2023 | 2D | 65,556,583 | 65.97 | 4.23 |
| Stomatal Area | Abaxial | TOS1 | 2023 | 3A | 648,742,426 | -46.30 | 2.86 |
| Stomatal Area | Abaxial | TOS1 | 2023 | 3B | 606,549,076 | -34.37 | 3.97 |
| Stomatal Area | Abaxial | TOS1 | 2023 | 3D | 36,703,071 | -55.94 | 7.57 |
| Stomatal Area | Adaxial | TOS1 | 2023 | 1A | 376,842,796 | 113.48 | 3.18 |

| | | | | | | | |
|---|---|---|---|---|---|---|---|
| Stomatal Area | Adaxial | TOS1 | 2023 | 1B | 555,412,597 | 72.26 | 5.73 |
| Stomatal Area | Adaxial | TOS1 | 2023 | 2B | 461,678,344 | -51.73 | 4.36 |
| Stomatal Area | Adaxial | TOS1 | 2023 | 5A | 665,150,454 | 76.71 | 4.36 |
| Stomatal Area | Adaxial | TOS1 | 2023 | 6B | 482,975,955 | -59.35 | 6.37 |
| Stomatal Area | Adaxial | TOS1 | 2023 | 7A | 23,804,088 | -37.38 | 4.12 |
| Stomatal Area | Adaxial | TOS1 | 2023 | 7A | 614,132,821 | 125.60 | 6.65 |
| Stomatal Area | Adaxial | TOS1 | 2023 | 7B | 457,582,679 | -44.25 | 5.94 |
| Stomatal Area | Abaxial | TOS2 | 2023 | 1B | 555,411,043 | -56.89 | 3.99 |
| Stomatal Area | Abaxial | TOS2 | 2023 | 3A | 648,742,426 | -80.48 | 5.50 |
| Stomatal Area | Abaxial | TOS2 | 2023 | 7D | 56,960,435 | 64.32 | 5.51 |
| Stomatal Area | Adaxial | TOS2 | 2023 | 3B | 132,315,022 | 71.24 | 9.10 |
| Stomatal Area | Adaxial | TOS2 | 2023 | 3B | 427,210,130 | -45.94 | 4.10 |
| Stomatal Area | Adaxial | TOS2 | 2023 | 5B | 461,435,203 | 54.50 | 9.51 |
| Stomatal Area | Adaxial | TOS2 | 2023 | 5B | 713,283,368 | 39.38 | 4.98 |
| Stomatal Area | Adaxial | TOS2 | 2023 | 7B | 225,050,548 | -52.63 | 3.76 |
| Stomatal Area | Abaxial | TOS1 | 2024 | 2B | 465,607,423 | -76.52 | 7.31 |
| Stomatal Area | Abaxial | TOS1 | 2024 | 4B | 543,140,936 | 50.02 | 3.83 |

| | | | | | | | |
|---|---|---|---|---|---|---|---|
| Stomatal Area | Adaxial | TOS1 | 2024 | 3B | 247,910,335 | 87.25 | 6.67 |
| Stomatal Area | Abaxial | TOS2 | 2024 | 5B | 406,738,159 | 53.82 | 2.62 |
| Stomatal Area | Abaxial | TOS2 | 2024 | 7A | 549,012,379 | 102.58 | 3.36 |
| Stomatal Area | Abaxial | TOS2 | 2024 | 7D | 6,846,653 | 86.27 | 5.23 |
| Stomatal Density | Abaxial | TOS1 | 2023 | 1B | 518,446,586 | 0.15 | 4.21 |
| Stomatal Density | Abaxial | TOS1 | 2023 | 5A | 592,814,280 | -0.11 | 5.10 |
| Stomatal Density | Abaxial | TOS1 | 2023 | 6A | 581,018,698 | 0.19 | 5.13 |
| Stomatal Density | Abaxial | TOS1 | 2023 | 7B | 547,215,666 | 0.17 | 7.04 |
| Stomatal Density | Adaxial | TOS1 | 2023 | 1A | 377,335,032 | 0.27 | 3.15 |
| Stomatal Density | Adaxial | TOS1 | 2023 | 1A | 574,778,628 | 0.13 | 3.58 |
| Stomatal Density | Adaxial | TOS1 | 2023 | 5A | 592,188,836 | 0.10 | 4.27 |
| Stomatal Density | Adaxial | TOS1 | 2023 | 7B | 526,910,362 | 0.20 | 5.92 |
| Stomatal Density | Abaxial | TOS2 | 2023 | 2B | 142,124,038 | -0.13 | 4.29 |
| Stomatal Density | Abaxial | TOS2 | 2023 | 7B | 526,913,571 | 0.13 | 3.44 |
| Stomatal Density | Adaxial | TOS2 | 2023 | 5B | 681,423,212 | 0.16 | 6.05 |
| Stomatal Density | Adaxial | TOS2 | 2023 | 5D | 552,685,132 | -0.17 | 5.35 |
| Stomatal Density | Abaxial | TOS1 | 2024 | 2A | 709,716,725 | 0.43 | 3.67 |
| $g_s$ | Abaxial | TOS1 | 2023 | 3A | 743,381,074 | 0.02 | 3.85 |

| | | | | | | | |
|---|---|---|---|---|---|---|---|
| $g_s$ | Abaxial | TOS2 | 2023 | 1A | 534,461,457 | -0.02 | 4.44 |
| $g_s$ | Abaxial | TOS2 | 2023 | 2A | 384,736,045 | 0.01 | 2.61 |
| $g_s$ | Abaxial | TOS2 | 2023 | 2B | 192,541,599 | 0.02 | 5.41 |
| $g_s$ | Abaxial | TOS2 | 2023 | 3D | 40,608,481 | -0.03 | 5.37 |
| $g_s$ | Abaxial | TOS2 | 2023 | 7A | 423,478,562 | 0.02 | 3.21 |
| $g_{smax}$ | Abaxial | TOS1 | 2023 | 1A | 15,656,623 | 0.07 | 3.67 |
| $g_{smax}$ | Abaxial | TOS1 | 2023 | 2B | 255,839,971 | 0.03 | 3.70 |
| $g_{smax}$ | Abaxial | TOS1 | 2023 | 2B | 789,702,543 | -0.09 | 8.51 |
| $g_{smax}$ | Abaxial | TOS1 | 2023 | 3B | 756,969,739 | -0.07 | 4.75 |
| $g_{smax}$ | Abaxial | TOS1 | 2023 | 6A | 581,018,698 | 0.07 | 6.83 |
| $g_{smax}$ | Abaxial | TOS1 | 2023 | 7B | 82,082,909 | -0.04 | 3.78 |
| $g_{smax}$ | Abaxial | TOS1 | 2023 | 7B | 547,215,666 | 0.06 | 7.00 |
| $g_{smax}$ | Adaxial | TOS1 | 2023 | 1A | 575,536,483 | 0.06 | 6.60 |
| $g_{smax}$ | Adaxial | TOS1 | 2023 | 1B | 579,462,182 | 0.04 | 5.78 |
| $g_{smax}$ | Adaxial | TOS1 | 2023 | 7A | 249,226,230 | -0.03 | 4.64 |
| $g_{smax}$ | Adaxial | TOS1 | 2023 | 7B | 62,016,354 | -0.06 | 4.68 |
| $g_{smax}$ | Adaxial | TOS1 | 2023 | 7B | 526,910,362 | 0.05 | 4.18 |
| $g_{smax}$ | Adaxial | TOS2 | 2023 | 4B | 534,200,348 | -0.03 | 3.44 |
| $g_{smax}$ | Adaxial | TOS2 | 2023 | 5B | 681,423,212 | 0.05 | 6.19 |
| $g_{smax}$ | Adaxial | TOS2 | 2023 | 7B | 62,057,627 | -0.06 | 4.86 |
| $g_{se}$ | Abaxial | TOS1 | 2023 | 3A | 743,381,074 | 0.02 | 3.02 |
| $g_{se}$ | Abaxial | TOS1 | 2023 | 4A | 459,394,052 | 0.03 | 3.15 |
| $g_{se}$ | Abaxial | TOS1 | 2023 | 6D | 484,299,874 | 0.02 | 4.19 |
| $g_{se}$ | Abaxial | TOS2 | 2023 | 1A | 519,184,553 | 0.02 | 3.67 |

**Table S6: Season 2 (2024) data showing average values for key traits at TOS 1 and at TOS 2, and % change from TOS 1 to TOS 2. Values are given for each surface independently and for both surfaces averaged.**

| Trait & Surface | | TOS 1 | TOS 2 | % Change TOS 1 to TOS 2 |
|---|---|---|---|---|
| **Abaxial** | | | | |
| $g_s$ | $mol.m^{-2}.s^{-1}$ | 0.171 | 0.137 | -19.89 |
| **Stomatal Density** | per $mm^{-2}$ | 35.38 | 40.16 | 13.52 |
| **Guard cell Length** | µm | 65.31 | 61.72 | -5.50 |
| **Guard Cell Width** | µm | 38.42 | 38.58 | 0.41 |
| **Stomatal Area** | $\mu m^{-2}$ | 1947.47 | 1855.96 | -4.70 |
| $g_{smax}$ | $mol.m^{-2}.s^{-1}$ | 0.97 | 1.04 | 7.14 |
| $g_{se}$ | - | 0.18 | 0.14 | -23.64 |
| **Adaxial** | | | | |
| $g_s$ | $mol.m^{-2}.s^{-1}$ | 0.347 | 0.374 | 7.90 |
| **Stomatal Density** | per $mm^{-2}$ | 46.94 | 50.36 | 7.29 |
| **Guard cell Length** | µm | 66.87 | 62.30 | -6.83 |
| **Guard Cell Width** | µm | 35.88 | 36.02 | 0.41 |
| **Stomatal Area** | $\mu m^{-2}$ | 1844.24 | 1730.59 | -6.16 |
| $g_{smax}$ | $mol.m^{-2}.s^{-1}$ | 1.32 | 1.32 | -0.28 |
| $g_{se}$ | - | 0.26 | 0.29 | 9.45 |
| **Adaxial & Abaxial Average** | | | | |
| $g_s$ | $mol.m^{-2}.s^{-1}$ | 0.259 | 0.256 | -1.27 |
| **Stomatal Density** | per $mm^{-2}$ | 41.16 | 45.26 | 9.97 |
| **Guard cell Length** | µm | 66.09 | 62.01 | -6.17 |
| **Guard Cell Width** | µm | 37.15 | 37.30 | 0.41 |
| **Stomatal Area** | $\mu m^{-2}$ | 1895.86 | 1793.27 | -5.41 |
| $g_{smax}$ | $mol.m^{-2}.s^{-1}$ | 1.15 | 1.18 | 2.87 |
| $g_{se}$ | - | 0.22 | 0.21 | -3.87 |
| **Yield** | $t.ha^{-1}$ | 5.88 | 3.48 | -40.75 |
| **Thousand Kernel Weight** | g | 41.16 | 34.96 | -15.06 |
| **Screenings** | % | 3.09 | 5.82 | 88.43 |
| **Moisture** | % | 10.48 | 9.86 | -5.87 |
| **Protein** | % | 12.05 | 12.84 | 6.54 |
| **Test Weight** | $kg.hL^{-1}$ | 83.82 | 79.77 | -4.83 |

**Table S7: The number of QTLs reported in literature by trait and chromosome. The last column shows the number of putative QTLs we found for the same trait and chromosome.**

| Trait | Chromosome | # Reported QTLs | Source | # Putative QTLs found in our study |
|---|---|---|---|---|
| SA | 1B | 3 | Shahinnia et al. 2016, Liu et al. 2025 | 3 |
| SA | 2B | 2 | Liu et al. 2025 | 2 |
| SA | 4A | 1 | Shahinnia et al. 2016 | 0 |
| SA | 4B | 5 | Shahinnia et al. 2016, Liu et al. 2025 | 1 |
| SA | 5A | 2 | Shahinnia et al. 2016, Liu et al. 2025 | 1 |
| SA | 5B | 15 | Shahinnia et al. 2016, Liu et al. 2025 | 3 |
| SA | 5D | 3 | Shahinnia et al. 2016, Liu et al. 2025 | 0 |
| SD | 1A | 3 | Liu et al. 2025 | 2 |
| SD | 3A | 5 | Shahinnia et al. 2016, Liu et al. 2025 | 0 |
| SD | 4A | 1 | Shahinnia et al. 2016 | 0 |
| SD | 4B | 2 | Shahinnia et al. 2016, Liu et al. 2025 | 0 |
| SD | 5A | 7 | Shahinnia et al. 2016, Liu et al. 2025 | 2 |
| SD | 5B | 2 | Shahinnia et al. 2016, Liu et al. 2025 | 1 |
| SD | 7A | 3 | Shahinnia et al. 2016, Liu et al. 2025 | 0 |
| $g_s$ | 2B | 1 | Wang et al. 2015 | 1 |
| $g_s$ | 2D | 1 | Wang et al. 2015 | 0 |
| $g_s$ | 4A | 1 | Wang et al. 2015 | 0 |
| $g_s$ | 6D | 1 | Wang et al. 2015 | 0 |
| $g_s$ | 7B | 2 | Wang et al. 2015 | 0 |

| | | | | |
|---|---|---|---|---|
| GCL | 1A | 1 | Shahinnia et al. 2016 | 0 |
| GCL | 1B | 5 | Shahinnia et al. 2016, Liu et al. 2025 | 1 |
| GCL | 3B | 1 | Shahinnia et al. 2016 | 1 |
| GCL | 4B | 8 | Shahinnia et al. 2016, Liu et al. 2025 | 2 |
| GCL | 5B | 3 | Liu et al. 2025 | 5 |
| GCL | 7A | 3 | Shahinnia et al. 2016 | 1 |
| GCL | 7D | 1 | Shahinnia et al. 2016 | 5 |
| GCW | 2A | 1 | Liu et al. 2025 | 0 |
| GCW | 2B | 1 | Liu et al. 2025 | 3 |
| GCW | 5B | 7 | Liu et al. 2025 | 3 |
| GCW | 5D | 1 | Liu et al. 2025 | 2 |
| GCW | 7A | 4 | Liu et al. 2025 | 3 |

**Table S8: The number of putative QTLs within a 10 Mbp region by chromosome and trait. The 20 bolded rows indicate a region that may contain pleiotropic QTLs for stomatal traits.**

| **Chromosome** | **Range of region (Mbp)** | **GCL** | **GCW** | **SA** | **SD** | $g_s$ | $g_{smax}$ | $g_{se}$ |
|---|---|---|---|---|---|---|---|---|
| 1A | (15,25] | 0 | 0 | 0 | 0 | 0 | 1 | 0 |
| **1A** | **(375,385]** | **0** | **1** | **1** | **1** | **0** | **0** | **0** |
| 1A | (515,525] | 0 | 0 | 0 | 0 | 0 | 0 | 1 |
| 1A | (525,535] | 0 | 0 | 0 | 0 | 1 | 0 | 0 |
| 1A | (545,555] | 0 | 1 | 0 | 0 | 0 | 0 | 0 |
| 1A | (565,575] | 0 | 0 | 0 | 1 | 0 | 0 | 0 |
| **1A** | **(575,585]** | **0** | **1** | **0** | **0** | **0** | **1** | **0** |
| 1B | (165,175] | 1 | 0 | 0 | 0 | 0 | 0 | 0 |
| 1B | (405,415] | 0 | 1 | 0 | 0 | 0 | 0 | 0 |
| 1B | (455,465] | 0 | 1 | 0 | 0 | 0 | 0 | 0 |
| 1B | (515,525] | 0 | 0 | 0 | 1 | 0 | 0 | 0 |
| 1B | (555,565] | 0 | 0 | 3 | 0 | 0 | 0 | 0 |
| 1B | (575,585] | 0 | 0 | 0 | 0 | 0 | 1 | 0 |
| 1B | (585,595] | 0 | 1 | 0 | 0 | 0 | 0 | 0 |
| 2A | (375,385] | 0 | 0 | 0 | 0 | 1 | 0 | 0 |
| 2A | (705,715] | 0 | 0 | 0 | 1 | 0 | 0 | 0 |
| 2B | [5,15] | 0 | 1 | 0 | 0 | 0 | 0 | 0 |
| 2B | (135,145] | 0 | 0 | 0 | 1 | 0 | 0 | 0 |
| 2B | (185,195] | 0 | 0 | 0 | 0 | 1 | 0 | 0 |
| 2B | (255,265] | 0 | 0 | 0 | 0 | 0 | 1 | 0 |
| 2B | (395,405] | 0 | 1 | 0 | 0 | 0 | 0 | 0 |
| **2B** | **(455,465]** | **0** | **1** | **1** | **0** | **0** | **0** | **0** |
| 2B | (465,475] | 0 | 0 | 1 | 0 | 0 | 0 | 0 |
| 2B | (655,665] | 1 | 0 | 0 | 0 | 0 | 0 | 0 |
| 2B | (685,695] | 1 | 0 | 0 | 0 | 0 | 0 | 0 |

| | | | | | | | | |
|---|---|---|---|---|---|---|---|---|
| 2B | (785,795] | 0 | 0 | 0 | 0 | 0 | 1 | 0 |
| 2D | (65,75] | 0 | 0 | 1 | 0 | 0 | 0 | 0 |
| 2D | (585,595] | 0 | 1 | 0 | 0 | 0 | 0 | 0 |
| 3A | (45,55] | 1 | 0 | 0 | 0 | 0 | 0 | 0 |
| 3A | (645,655] | 0 | 0 | 2 | 0 | 0 | 0 | 0 |
| **3A** | **(735,745]** | **1** | **0** | **0** | **0** | **1** | **0** | **1** |
| 3B | (45,55] | 0 | 1 | 0 | 0 | 0 | 0 | 0 |
| 3B | (65,75] | 1 | 0 | 0 | 0 | 0 | 0 | 0 |
| 3B | (125,135] | 0 | 0 | 1 | 0 | 0 | 0 | 0 |
| 3B | (245,255] | 0 | 0 | 1 | 0 | 0 | 0 | 0 |
| 3B | (425,435] | 0 | 0 | 1 | 0 | 0 | 0 | 0 |
| 3B | (555,565] | 0 | 1 | 0 | 0 | 0 | 0 | 0 |
| 3B | (605,615] | 0 | 0 | 1 | 0 | 0 | 0 | 0 |
| 3B | (755,765] | 0 | 0 | 0 | 0 | 0 | 1 | 0 |
| **3D** | **(35,45]** | **0** | **0** | **1** | **0** | **1** | **0** | **0** |
| 4A | (115,125] | 0 | 1 | 0 | 0 | 0 | 0 | 0 |
| 4A | (455,465] | 0 | 0 | 0 | 0 | 0 | 0 | 1 |
| 4B | (25,35] | 1 | 0 | 0 | 0 | 0 | 0 | 0 |
| 4B | (525,535] | 0 | 0 | 0 | 0 | 0 | 1 | 0 |
| 4B | (535,545] | 0 | 0 | 1 | 0 | 0 | 0 | 0 |
| 4B | (655,665] | 1 | 0 | 0 | 0 | 0 | 0 | 0 |
| 5A | (545,555] | 1 | 0 | 0 | 0 | 0 | 0 | 0 |
| **5A** | **(585,595]** | **2** | **0** | **0** | **2** | **0** | **0** | **0** |
| **5A** | **(665,675]** | **1** | **0** | **1** | **0** | **0** | **0** | **0** |
| 5B | (55,65] | 0 | 1 | 0 | 0 | 0 | 0 | 0 |
| **5B** | **(405,415]** | **3** | **0** | **1** | **0** | **0** | **0** | **0** |
| **5B** | **(455,465]** | **1** | **0** | **1** | **0** | **0** | **0** | **0** |
| 5B | (525,535] | 0 | 1 | 0 | 0 | 0 | 0 | 0 |
| 5B | (545,555] | 1 | 0 | 0 | 0 | 0 | 0 | 0 |

| | | | | | | | | |
|---|---|---|---|---|---|---|---|---|
| **5B** | **(675,685]** | **0** | **0** | **0** | **1** | **0** | **1** | **0** |
| **5B** | **(705,715]** | **0** | **1** | **1** | **0** | **0** | **0** | **0** |
| **5D** | **(545,555]** | **0** | **2** | **0** | **1** | **0** | **0** | **0** |
| **6A** | **(575,585]** | **0** | **0** | **0** | **1** | **0** | **1** | **0** |
| 6A | (605,615] | 1 | 0 | 0 | 0 | 0 | 0 | 0 |
| 6B | (15,25] | 0 | 2 | 0 | 0 | 0 | 0 | 0 |
| **6B** | **(475,485]** | **1** | **1** | **1** | **0** | **0** | **0** | **0** |
| 6B | (605,615] | 1 | 0 | 0 | 0 | 0 | 0 | 0 |
| 6B | (715,725] | 0 | 1 | 0 | 0 | 0 | 0 | 0 |
| 6B | (725,735] | 0 | 1 | 0 | 0 | 0 | 0 | 0 |
| 6D | (475,485] | 0 | 0 | 0 | 0 | 0 | 0 | 1 |
| 7A | (15,25] | 0 | 0 | 1 | 0 | 0 | 0 | 0 |
| 7A | (175,185] | 1 | 0 | 0 | 0 | 0 | 0 | 0 |
| 7A | (245,255] | 0 | 0 | 0 | 0 | 0 | 1 | 0 |
| 7A | (415,425] | 0 | 0 | 0 | 0 | 1 | 0 | 0 |
| 7A | (545,555] | 0 | 0 | 1 | 0 | 0 | 0 | 0 |
| **7A** | **(605,615]** | **0** | **1** | **1** | **0** | **0** | **0** | **0** |
| 7A | (675,685] | 0 | 2 | 0 | 0 | 0 | 0 | 0 |
| 7B | (55,65] | 0 | 0 | 0 | 0 | 0 | 2 | 0 |
| 7B | (75,85] | 0 | 0 | 0 | 0 | 0 | 1 | 0 |
| 7B | (145,155] | 0 | 1 | 0 | 0 | 0 | 0 | 0 |
| 7B | (225,235] | 0 | 0 | 1 | 0 | 0 | 0 | 0 |
| **7B** | **(455,465]** | **2** | **0** | **1** | **0** | **0** | **0** | **0** |
| **7B** | **(525,535]** | **1** | **0** | **0** | **2** | **0** | **1** | **0** |
| **7B** | **(545,555]** | **0** | **0** | **0** | **1** | **0** | **1** | **0** |
| 7B | (695,705] | 0 | 1 | 0 | 0 | 0 | 0 | 0 |
| 7B | (715,725] | 0 | 1 | 0 | 0 | 0 | 0 | 0 |
| 7B | (725,735] | 0 | 1 | 0 | 0 | 0 | 0 | 0 |
| **7D** | **[5,15]** | **1** | **0** | **1** | **0** | **0** | **0** | **0** |

| | | | | | | | | |
|---|---|---|---|---|---|---|---|---|
| **7D** | **(55,65]** | **4** | **1** | **1** | **0** | **0** | **0** | **0** |
